# Fast and parallel nanoscale 3D tracking of heterogeneous mammalian chromatin dynamics

**DOI:** 10.1101/2021.10.24.465625

**Authors:** Anna-Karin Gustavsson, Rajarshi P. Ghosh, Petar N. Petrov, Jan T. Liphardt, W. E. Moerner

## Abstract

Chromatin organization and dynamics are critical for gene regulation. In this work we present a methodology for fast and parallel 3D tracking of multiple chromosomal loci of choice over many thousands of frames on various time scales. We achieved this by developing and combining fluorogenic and replenishable nanobody arrays, engineered point spread functions, and light sheet illumination. The result is gentle live-cell 3D tracking with excellent spatiotemporal resolution throughout the mammalian cell nucleus. Correction for both sample drift and nuclear translation facilitated accurate long-term tracking of the chromatin dynamics. We demonstrate tracking of both fast dynamics (50 Hz) and over time scales extending to several hours, and we find both large heterogeneity between cells and apparent anisotropy in the dynamics in the axial direction. We further quantify the effect of inhibiting actin polymerization on the dynamics and find an overall increase in both the apparent diffusion coefficient D* and anomalous diffusion exponent α, and a transition to more isotropic dynamics in 3D after such treatment. We think that our methodology in the future will allow researchers to obtain a better fundamental understanding of chromatin dynamics and how it is altered during disease progression and after perturbations of cellular function.

## INTRODUCTION

The human genome is highly organized within the cell nucleus and this dynamic 3D architecture plays a critical role in gene regulation and activity^1^. Over the last decades, major advances have been made in our understanding of the chromatin organization using sequencing-based techniques such as Hi-C^2^ and other DNA-DNA proximity methods^3^. However, to fully understand how the genome operates requires complementary methods that can access spatial information about 3D chromatin dynamics and genome reorganization in individual living cells. Fluorescent labeling and tracking of chromosomal loci is a method that allows extraction of such dynamic information. However, several crucial factors have limited previous methods for long term, fast, parallel 3D tracking of multiple chromosomal loci.

First, loci of choice should be labeled with a fluorescent tag that is bright and photostable, and that yields low cellular fluorescence background. Tagging of loci using single fluorescent proteins fused to deactivated (d)Cas9 allows for targeting to specific chromosomal loci of choice using guide RNAs^4^. However, this method is limited by low signal combined with short track lengths and high fluorescence background from unbound fluorophores. One solution is to fuse or recruit multiple fluorescent proteins to a target, as has been demonstrated by approaches such as the SunTag^5^, FP11-tags^6^, MS2 tags^7^, PP7 tags^8^, and Spaghetti monster^9^. However, these options can suffer from high background and/or slow maturation. Recently, we developed a method utilizing a nuclear-importable array of the enhancer GFP nanobody GBP1^10^, called ArrayG/N, to overcome the issues with previous labeling schemes^11^. Using this approach, we achieved temporally unlimited 2D tracking of kinesins, integrins, and histones when combined with TIRF illumination. However, up until now this approach has not been demonstrated for direct labeling and tracking of chromosomal loci.

Tracking of chromosomal loci has most conventionally been performed using wide-field epi-illumination, TIRF, or confocal imaging, often using maximum projections of multiple z-slices. However, chromatin dynamics is not necessarily isotropic or confined to a single 2D plane within the nucleus, and only extracting 2D information from an inherently 3D system can lead to misinterpretation of the dynamics. Conventional confocal imaging in particular only provides 3D information after multiple slices are acquired, which can miss motions which occur in slices not being imaged at a given moment. To extract information about fast and potentially anisotropic dynamics and to facilitate long-term tracking of multiple unique loci without losing them out of the detection volume requires wide-field 3D detection over an extended axial range. Fast 3D tracking over an extended axial range has been demonstrated using confocal active-feedback approaches, such as orbital imaging^12,13^ and TSUNAMI^14^. However, these methods have been limited to tracking of a maximum of two loci simultaneously. Biplane^15–17^ and multiplane^18–22^ imaging are wide-field approaches that have been implemented for tracking inside the nucleus, but these methods can be restricted by either limited axial range or by low temporal resolution by weaker signals due to splitting of the light into multiple planes. An alternative approach we show here is to use engineered point spread functions (PSFs) (for a review, see ref ^23^), where the axial (z) position of the emitter is encoded directly in the shape of the PSF on the camera. This is accomplished by applying a phase pattern to the emitted light in the Fourier plane of the microscope and allows for scan-free wide-field 3D detection of emitters with excellent precision by the addition of a small number of optical elements to the standard fluorescence microscope. This method has utilized multiple different types of PSFs with various axial ranges between 1-20 μm, including the astigmatic^24–28^, bisected pupil^29^, self-bending^30^, corkscrew^31^, double helix (DH)^32–35^, and Tetrapod PSFs^36–39^.

The spatiotemporal resolution that can be achieved for 3D tracking using engineered PSFs depends on the signal-to-background ratio (SBR) between the fluorescence signal from the locus and the background fluorescence from the rest of the cell. Live-cell tracking also requires gentle illumination to reduce the risk of photodamage. Light sheet illumination, where the sample is optically sectioned by illumination with a thin sheet of light introduced from the side, is a wide-field illumination approach that allows for good contrast and gentle live-cell imaging and that is compatible with tracking away from the coverslip^40,41^. Several light sheet designs have been developed to improve single-molecule imaging and single-particle tracking (for a review, see ref^42^). We recently developed a tilted light sheet design that alleviates drawbacks of previous designs and improves the resolution for 3D single-molecule imaging in fixed cells^32^.

In this work, we have developed a methodology that allows for parallel tracking of multiple chromosomal loci in 3D throughout the nucleus over many thousands of frames on various time scales. We have accomplished this by developing and combining (i) nanobody array labeling of chromosomal loci based on the recently developed ArrayG/N scheme^11^ to achieve very bright loci, low fluorescence background, and replenishment of fluorophores to the loci over time; (ii) engineered PSFs for parallel 3D tracking of multiple chromosomal loci with a localization precision of tens of nm and temporal resolution up to 50 Hz; and (iii) tilted light sheet illumination to further reduce background fluorescence, reduce photodamage for live-cell tracking, and reduce photobleaching of fluorophores which further increases the achievable track lengths when combined with our replenishable labeling scheme. Chromosomal loci tracking was combined with tracking of fiducial beads melted to the coverslip using a long-range PSF and labeling and tracking of DNA to facilitate correction of sample drift and nuclear translation. We demonstrate our approach for tracking of both fast dynamics (50 Hz) and also over time scales ranging up to several hours. By fitting the dynamics to a fractional Brownian diffusion model, we extract dynamic parameters for both ensemble and individual tracks independently in x, y, and z, finding both large heterogeneity between cells and between loci in the same cell, as well as apparent anisotropy in the dynamics in the axial direction. We further quantify the effect of inhibiting actin polymerization on the dynamics and find an overall increase in both the apparent diffusion coefficient D* and the anomalous diffusion exponent α, and a transition to more homogenous dynamics in 3D after such treatment. Our methodology is flexible and can be targeted to chromatin regions of choice to determine the effect of a wide range of cell conditions and drug treatments on the chromatin dynamics.

## RESULTS

### Nanobody array labeling scheme for bright loci, low background, and fluorophore replenishment over time

Labeling was based on the ArrayG/N scheme using arrays of 16 GFP nanobodies (ArrayG/N)_16X_^11^ fused to the nuclease-deficient version of *Streptococcus pyogenes* Cas9 (dSpCas9) (Figure 1A). The arrays were then targeted to the desired chromosomal loci using single guide RNA (sgRNA) and labeled by monomeric wild-type GFP (mwGFP)^11^. Multiple binding sites per array results in very bright loci. Using dCas9 and sgRNA to target the chromosomal loci allows the researcher to conveniently select and change the chromosomal loci of choice for study. The choice of mwGFP as fluorescent label has several benefits over the more conventional enhanced (e)GFP: first, mwGFP did not show clustering, as was the case when using eGFP^11^, and second, mwGFP has a fluorogenic behavior when used together with the selected GFP nanobodies. Specifically, mwGFP is initially dim when unbound in the cell, but it undergoes significant fluorescence enhancement of ∼26-fold upon binding to the array^11^. This effect drastically reduces the fluorescence background while maintaining high signal at the loci, and the resulting excellent contrast improves our localization precision when tracking the loci. Having a high SBR also allows us to reduce the laser intensity and/or the exposure time, which results in gentler live-cell tracking, longer tracks, and/or faster tracking. In addition, the binding to the array is reversible, which means that fluorophores bound to the array that photobleach can be replaced by unbleached fluorophores from the diffusing pool of fluorophores in the cell. This property is particularly beneficial when paired with light sheet illumination, where only a thin section around the tracked loci is illuminated, leaving the rest of the fluorophore pool intact.

**Figure 1.**
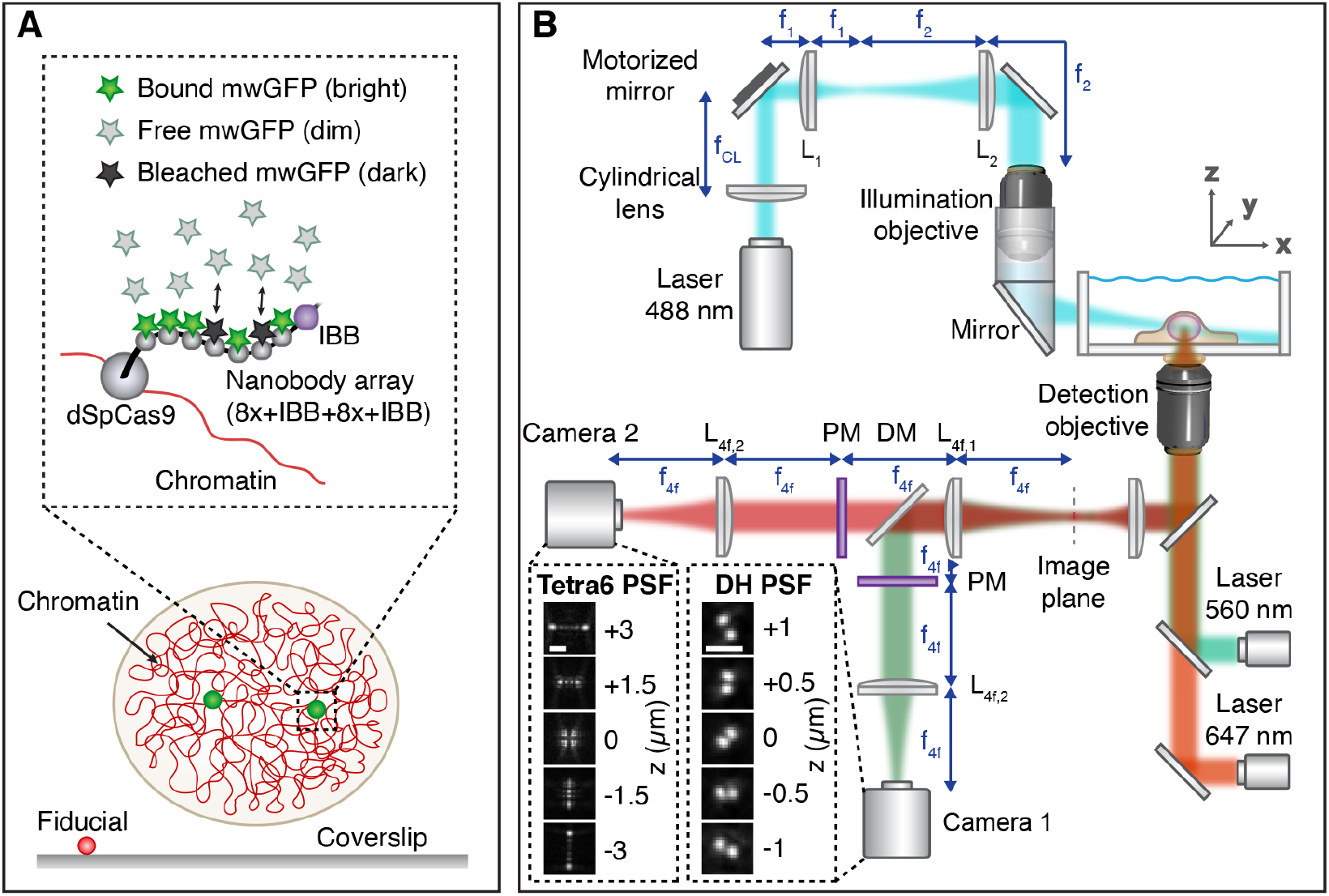
Experimental design. (A) Schematic showing the sample labeling scheme. Chromosomal loci were labeled using dSpCas9 fused to an array of 16 GFP nanobodies to which monomeric wildtype GFP (mwGFP) expressed by the cell could reversibly bind. Two Snurportin Importin Beta binding domains (IBB) were included to facilitate nuclear import. Chromatin was also labeled using SiR-DNA to allow tracking of nuclear translation. Fluorescent fiducial beads were melted to the coverslip to provide a measure of sample drift. (B) Simplified schematic of the experimental setup. Labeled chromosomal loci were excited by a tilted light sheet introduced from the side at an angle. The 3D positions of the loci were detected by imaging the emitted fluorescence light using the double-helix (DH) PSF with 2 µm axial range (Camera 1). Fiducial beads and chromatin were excited by epi-illumination at 560 nm and 647 nm, respectively, and emitted light from both the fiducial bead and the chromatin was detected using the Tetrapod PSF with 6 µm axial range (Camera 2). DM: dichroic mirror; PM: phase mask. Scale bars in the PSF images are 2 µm.

### Engineered PSFs for parallel 3D tracking with excellent spatiotemporal resolution

A DH PSF with a 2 µm axial range was used for tracking of chromosomal loci in a green channel, while a Tetrapod PSF with 6 µm axial range was used for tracking of fiducial beads in a red channel to facilitate drift correction in 3D (Figure 1B and Supplemental Figure S1). Using a shorter-range PSF for tracking of the loci allowed for tracking with very high precision, whereas using a longer-range PSF for tracking of fiducial beads allowed for detection at the coverslip even when positioning the image plane higher up in the sample. In the case of fiducial beads, the localization precision can be improved by tuning the intensity of the excitation light. Slight melting of the beads to the coverslip was required to avoid having the beads be endocytosed by the cells or becoming adhered to the cell surface. Any motion of the beads in relation to the coverslip would otherwise become convolved with the motion of the loci. For long-term experiments, the motion of the nucleus was also tracked using the Tetrapod PSF in the red channel by labeling DNA using SiR-DNA to allow for correction of nuclear translation. On shorter time scales nuclear translation was negligible compared to the motion of the loci. This approach typically provided median localization precision as small as 2-3 nm in xy and 4 nm in z for tracking at 0.5 Hz, and 7-8 nm in xy and 11 nm in z for tracking at 50 Hz. Distributions of localization precisions are provided in Supplemental Figure S2.

### Tilted light sheet illumination for gentler live cell tracking with improved precision and track lengths

A tilted light sheet was used to optically section the cells to further reduce fluorescence background, photobleaching, and the risk of photodamaging the live cells (Figure 1B). The tilted light sheet design was similar to the TILT3D design described previously^32^, but in this work the light sheet was introduced into the sample using a mirror mounted on an aluminum tube attached to the illumination objective. This approach decoupled any motion of the sample and the light sheet, which allowed for easy alignment of the light sheet with the image plane and easy scanning of the sample in x, y, and z. The mwGFP-labeled loci were excited by a 488 nm tilted light sheet that was ∼2.5 μm thick (1/e^2^), which improved the contrast by more than a factor of 2.5 compared to using conventional epi-illumination for tracking of the loci (Supplemental Figure S3). Using light sheet illumination in combination with our nanobody array labeling scheme is particularly advantageous due the reduced photobleaching of the unbound fluorophores, which allows for replenishment of fluorophores to the array.

Fiducial beads and DNA labeled with SiR-DNA were observed using epi-illumination at 560 nm and 647 nm, respectively. This approach allowed excitation of the beads and of the cell nucleus regardless of the position of the image plane and of the light sheet, and it also allowed for independent tuning of the illumination intensities for the three different lasers to ensure optimum precision without risking saturation in any channel.

### Correction for sample drift and nuclear translation

To accurately study the dynamics of the chromosomal loci within the cell nucleus, any sample drift must be removed in x, y, and z. For longer-term tracking, nuclear translation due to cell migration must also be removed, or it will become convolved with the measured motion of the loci. Our methodology to correct for sample drift and nuclear translation is described in more detail in Methods and is outlined in Supplemental Figure S1. To validate our approach, a labeled chromosomal locus in a fixed cell was tracked in the green channel using the DH PSF and the labeled nucleus and a fiducial bead were tracked in the red channel using the Tetrapod PSF while the microscope piezo stage was moved 20 nm each in x and y between frames to simulate nuclear translation (Figure 2, A and B). The resulting data show that all measured motions correspond well to the known stage motion, with resulting residuals over the 2000 nm total stage movement in x and y of [-33.6 ± 22.4, 5.7 ± 18.4, 10.2 ± 49.7] nm in x, y, and z, respectively, reported as mean ± standard deviation. The offset of the positions of the locus and the fiducial bead from the known stage position is consistent with stage drift during acquisition and shows the importance of proper drift correction. The importance of correcting for both sample drift and nuclear translation when tracking over longer time scales is demonstrated in Figure 2, C and D, where a locus track from a live cell is shown before and after correction.

**Figure 2.**
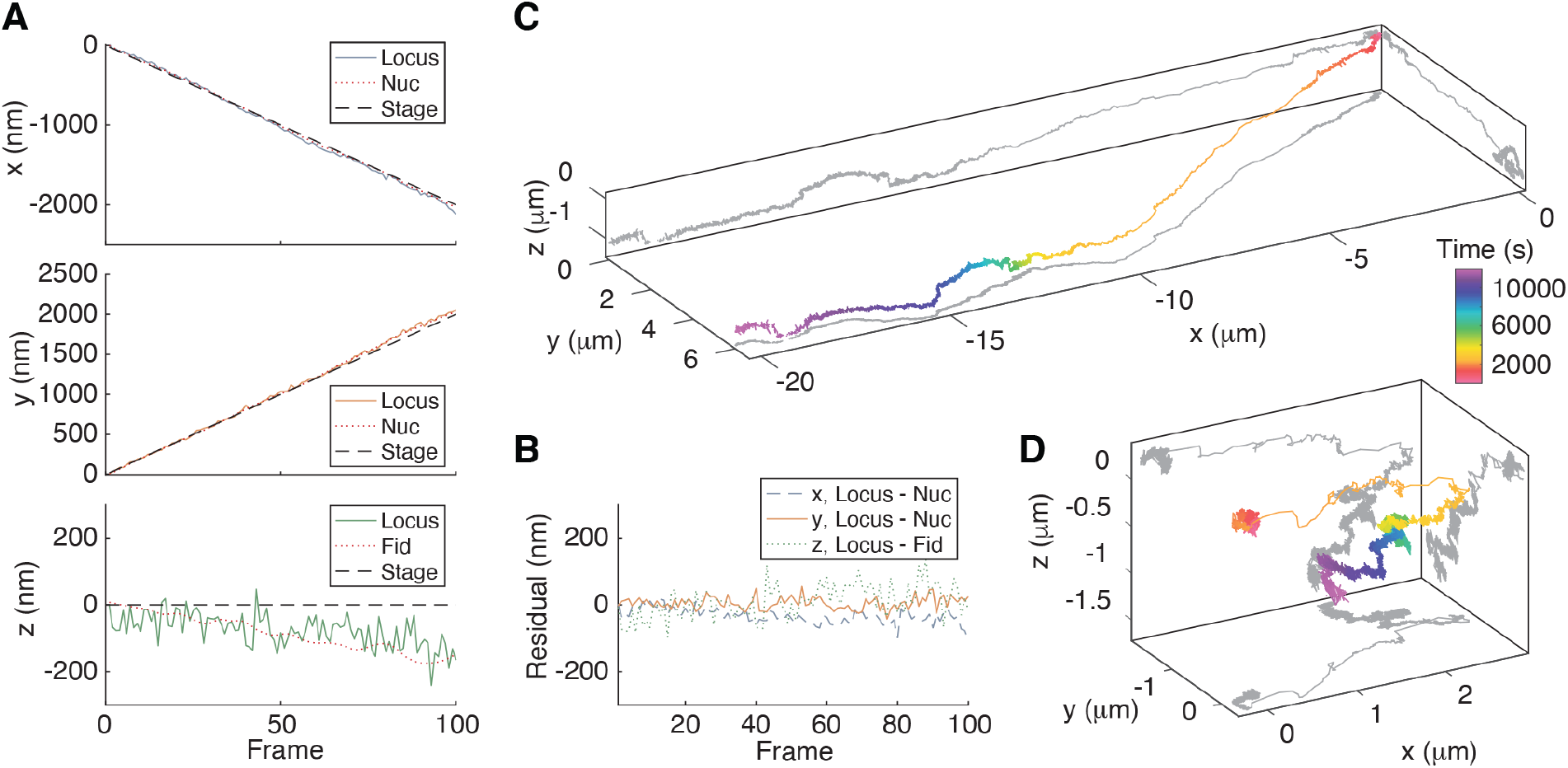
Validation of correction of sample drift and nuclear translation. (A) The labeled chromosomal locus in a fixed cell was tracked in the green channel using the DH PSF, and the labeled nucleus and a fiducial bead were tracked in the red channel using the Tetrapod PSF while the microscope piezo stage was moved 20 nm each in x and y between each frame to simulate nuclear translation. Sample drift was naturally occurring as during actual cell imaging. The measured movements in the red channel of the nucleus and of the fiducial bead were transformed into green coordinates, smoothed, and used to correct the measured motion of the locus. (B) Residual motion of the chromosomal locus after correction. The noisy appearance of the tracks is caused by localization imprecision of the relatively dim, fixed locus. (C) Apparent motion of a chromosomal locus in a live cell tracked at 0.5 Hz before any drift correction. (D) Motion of the locus in (C) after correction of sample drift and nuclear translation.

### Detailed chromatin dynamics in 3D at various time scales

To demonstrate the versatility of our methodology, parallel tracking of multiple chromosomal loci in live cells was performed at fast time scales (50 Hz, Figure 3A) and over longer time scales using time-lapse illumination and recording (0.5 Hz, Figure 3B). This resulted in track lengths of up to 10,000 frames, translating to multiple minutes at fast time scales and multiple hours for 0.5 Hz time-lapsing. The wide-field detection and extended axial range of the DH PSF allowed us to track multiple, unique loci in parallel over these time scales without having them diffuse out of range, which would have been the case for conventional 2D imaging with its limited depth-of-focus. Each track contains rich information about the local chromatin dynamics. In the example of longer time-scale tracking shown in Figure 3B, a large reorganization of the chromatin was detected after about 3000 s. This is an example of chromatin dynamics that our methodology will enable a better understanding of in the future.

**Figure 3.**
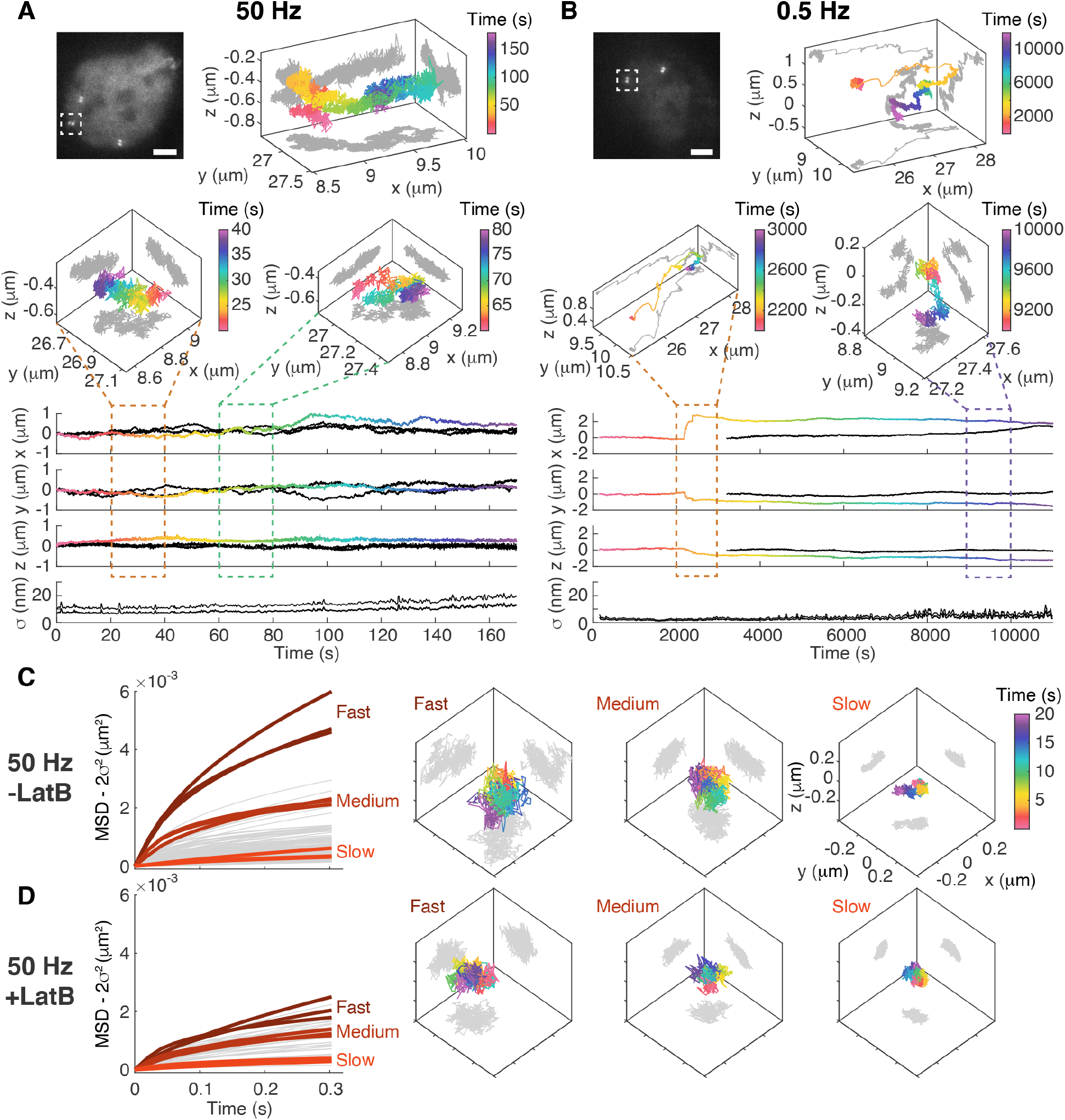
Very long 3D tracks of chromatin loci dynamics in live mammalian cells at short and long time scales. (A) The top left image shows a nucleus with four labeled loci of which three are in range for imaging with the DH PSF. The top 3D graph shows the resulting track from the locus in the white rectangle where images were acquired at 50 Hz. The bottom four graphs show the x, y, and z motion of the locus in the white rectangle (time color coded as in the 3D graph) as well as the motion of the other two loci (black), and the average localization precision *σ* of the locus in the white rectangle over a 15 data point running window. The 3D graph insets show the 3D dynamics in the marked time intervals. (B) The same type of data as in (A) but for a nucleus with two labelled loci imaged at 0.5 Hz. The second locus (black) came into range for imaging after about 3000 s. Scale bars are 5 µm. (C)-(D) Fitted MSDs from individual tracks and representative example tracks showing fast, medium, and slow dynamics for (C) untreated cells (-LatB) and (D) cells treated with latrunculin B (+LatB). All tracks are plotted on the same spatial and temporal scale. On average, LatB-treated cells showed faster dynamics.

Next, we set out to investigate the role of the actomyosin network on the chromatin 3D dynamics by inhibiting actin polymerization using latrunculin B (LatB). Quantifying the fast dynamics (50 Hz) of individual tracks in untreated live cells and cells treated with LatB by calculating the mean squared displacements (MSDs) revealed large heterogeneity in dynamics (Figure 3, C and D). Here, we show examples of tracks at three different time scales for each condition. In untreated cells, the majority of the loci showed relatively slow dynamics, but a few loci exhibited very fast dynamics. In cells treated with LatB, the average dynamical time scale was shifted toward shorter times. However, also here there is large heterogeneity in dynamics between different cells and between loci in the same cell.

### The 3D dynamics are anisotropic and affected by inhibition of actin polymerization

To further quantify the dynamics in untreated and LatB-treated cells, the MSDs were calculated for fast acquisitions (50 Hz) and fit using an expression (based on fractional Brownian motion as a source for anomalous diffusion) that takes errors arising from both localization precision and motion blur into account^43^. Extracting the ensemble average values of the effective diffusion coefficient D* and anomalous diffusion exponent α from live, untreated cells revealed an apparent anisotropy in the z-direction for both D* and α (Figure 4 and Supplemental Figure S4, and comparison with fixed cells in Supplemental Figure S5). As a control, the same experimental and analysis strategy when applied to 100 nm fluorescent polystyrene beads diffusing in a glycerol-water mixture resulted in isotropic, Brownian motion with an α value close to 1 that is expected from freely diffusing beads (Supplemental Figure S6). The measured anisotropy for live, untreated cells was also independent of the z-correction factor used to account for the index mismatch between the cell and the coverslip (Supplemental Table S1)^44^. To estimate what distribution of parameters can be expected from comparable numbers of tracks with varying track lengths and varying initial conditions of the fitting, we simulated 3D tracks of fractional Brownian motion (Supplemental Figures S7-S9). This showed as expected that a reduction of the number of localizations lead to a wider distribution, but that the analyses of the ensemble average results were robust with respect to the track lengths and initial conditions used for the experimental data.

**Figure 4.**
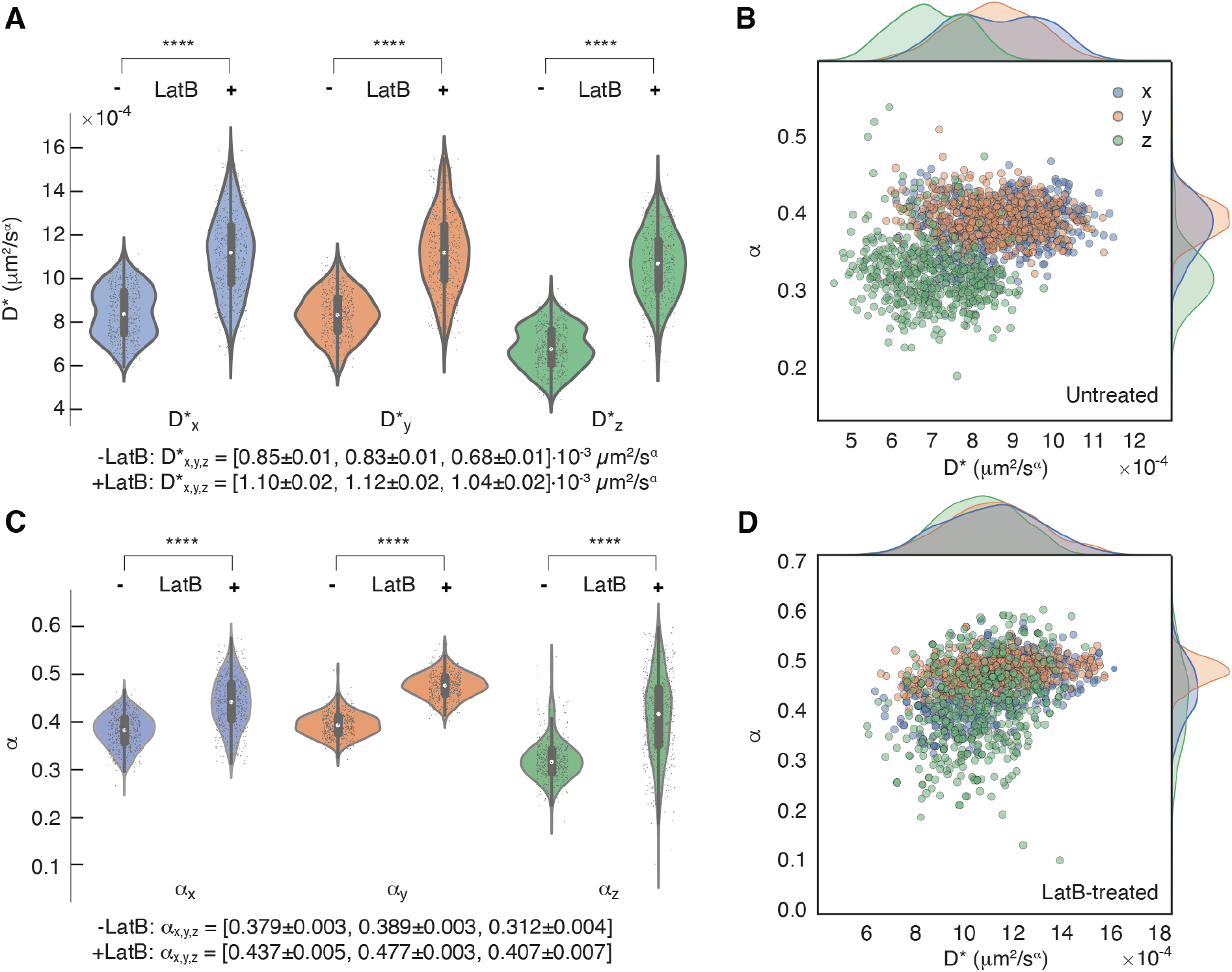
Analysis of 3D dynamics in untreated cells and cells treated with latrunculin B. The parameters were extracted from fits of the time-averaged ensemble MSDs using 500 bootstraps with 50% dropout in x, y, and z for untreated cells (-LatB) and cell treated with latrunculin B (+LatB). The complete data consisted of 54 tracks for untreated cells and 14 tracks for LatB-treated cells, each with a minimum track length of 640 frames. (A) Violin plots showing the resulting effective diffusion coefficient D*. The resulting values are reported as mean ± SEM. (B) Scatter plot and histograms showing the distribution of D* and α of untreated cells. (C) Violin plots showing the resulting α. The resulting values are reported as mean ± SEM. (D) Scatter plot and histograms showing the distribution of D* and α of LatB-treated cells. **** denotes *p* < 0.0001.

Inhibition of actin polymerization resulted in an overall increase in both D* and α, a wider distribution of these parameters, and a transition to more isotropic dynamics in 3D (Figure 4 and Supplemental Figure S10).

## DISCUSSION

The methodology developed in this work brings together the three different innovations of fluorogenic and replenishable nanobody array labeling, PSF engineering for 3D simultaneous tracking of multiple loci with excellent spatiotemporal resolution, and light sheet illumination for improved contrast, reduced photobleaching, and reduced photodamage of the live cells. This approach allowed us to track multiple chromosomal loci in parallel in 3D throughout the nucleus at fast (50 Hz) and slow (0.5 Hz) time scales for many thousands of frames with a precision of tens of nm while correcting for both sample drift and cell migration. Our results are consistent with previous data on lateral chromatin dynamics at short time scales^45,12,46–52,11^. However, by extracting dynamic parameters independently in x, y, and z for the ensemble and for individual tracks, we detect both an apparent axial anisotropy and large heterogeneity between cells and between different loci in the same cell.

The methodology developed in this work allowed us to investigate the role of the actomyosin cytoskeleton on chromatin dynamics and demonstrated how inhibition of latrunculin B leads to an overall increase in D* and α, similar to treatments with the actin polymerization inhibitor Cytochalasin D^53^. Owing to our 3D tracking, we also detected a transition to more isotropic motion after such treatment, which may be caused by a change in nuclear shape when forces on the nucleus by the actin cytoskeleton are relaxed.

Our method is flexible and yields nanoscale information about 3D chromatin dynamics on various time scales. We think that our approach in the future will allow us and other researchers to obtain a better fundamental understanding of chromatin dynamics and how the motions are affected during disease progression and after perturbations of cellular function.

## MATERIALS AND METHODS

### Array plasmid and guide plasmid generation

(ArrayG/N)_16X_ fusion of the nuclease-deficient version of *Streptococcus pyogenes* Cas9 (dSpCas9) was synthesized by concatenating dSpCas9 and two tandem repeats of (ArrayG/N)_8X_ in frame, where each unit of (ArrayG/N)_8X_ was composed of 8 copies of the codon optimized ArrayG unit^11^ and one copy of the snurportin importin beta binding domain (IBB). In linear sequence the fusion has the following plan: [(dSpCas9)-(40x glycine/serine linker)-(ArrayG)_8x_-(IBB)-(ArrayG)_8x_-(IBB)]. dSpCas9 and (ArrayG/N)_16X_ were ligated in frame into the multiple cloning site (MCS) of a custom-generated PiggyBac Tet-On 3G plasmid, where the cumate operator sequence (CuO) and EF1-CymR repressor-T2A-puromycin cassette of a PiggyBac cumate switch plasmid (PBQM812A-1, SBI) were replaced with a Tet-On 3G tetracycline inducible promoter and EF1-Tet-Activator-P2A-hygromycin cassette, respectively. This modification allowed leaky expression of the dSpCas9-(ArrayG/N)_16X_ fusion protein in the absence of doxycycline and high expression in the presence of doxycycline. To synthesize the binder plasmid, monomeric wild-type GFP (mwGFP)^11^ was inserted into the MCS of a PiggyBac cumate switch plasmid (PBQM812A-1, SBI), so that high expression of the binder could be achieved only after addition of the cumate inducer (QM100A-1, System Biosciences). (ArrayG/N)_16x_ has a total size of 255.6 kDa and of 689.2 kDa when fully occupied with mwGFP. The binding affinity of the nanobodies is 0.59 ± 0.11 nM^10^, and close to full occupancy is expected^11^.

For specific genomic site detection and tracking, sgRNA against the following repeated sequence (accactgtgatatcatacag) in chromosome 3 (interspersed in the genomic region spanning 195194259 to 195238776) was expressed from a U6 expression cassette in a lentiviral vector co-expressing blasticidin for antibiotic resistance from an SV40 early promoter.

### Cell line generation

To generate a stable cell line co-expressing (ArrayG/N)_16X_, mwGFP, and sgRNA, U2OS cells were first transfected with PiggyBac Tet-On 3G-(ArrayG/N)_16X_ and PBQM-mwGFP plasmids using the Neon Transfection system (Thermo Fisher Scientific) and the following pulse program (pulse voltage: 1050V, pulse width: 30 ms, 2 pulses). One day after transfection the growth medium was replaced with fresh medium. After three days of transfection, fresh growth medium supplemented with 1 µg/ml puromycin and 250 µg/ml hygromycin was added to the cells to initiate antibiotic selection. For the rest of the selection process antibiotic supplemented fresh growth medium was added to the cells every alternate day. The dual stable cell line was then transduced with lentivirus containing the sgRNA expression vector to generate the triple stable cell line. Briefly, viral particles produced by transfecting LentiX-293T (Takara Bio) cells with the sgRNA lentiviral vector were added to the dual stable cell lines. 3 days post viral transduction, fresh medium supplemented with 10 µg/ml blasticidin was added to initiate antibiotic selection.

### Cell culture

U2OS cells stably expressing (ArrayG/N)_16X_, mwGFP, and sgRNA were cultured at 37°C and 5% CO_2_ in high-glucose Dulbecco’s modified Eagle’s medium (DMEM, HyClone) supplemented with 10% (v/v) FBS (ClonTech), 250 µg/ml hygromycin, 1 µg/ml puromycin, and 10 µg/ml blasticidin. Expression of (ArrayG/N)_16X_ was induced by 1x doxycycline (1 µg/ml), and mwGFP was induced by 1x cumate (QM100A-1, System Biosciences) 24-48h before imaging. sgRNA was constitutively expressed.

### Sample preparation

Red fluorescent beads (580/605 nm, F8801, Invitrogen) diluted 1:100000 in nanopure water were spun onto plasma-etched coverslips (no. 1.5, 22×22 mm, Fisher Premium Cover Glass) at 10000 rpm for 30 s. The coverslips were then heated on a hotplate at 110°C for 20 min to attach the beads to the coverslip by gentle melting of the polystyrene. After cooling, the coverslips were fibronectin-coated (2 µl/cm^2^, Calbiochem, 341635) for 1h and washed 3x with PBS and 1x with cell culture medium. Cells were seeded at 25-30% confluency onto the prepared coverslips placed in 6-well plates in cell imaging media comprising phenol-red free DMEM (HyClone) supplemented with 10% (v/v) FBS (ClonTech) and 1 mM sodium pyruvate and allowed to settle for at least 4h. DNA was stained by adding 330 nM SiR-DNA (Spirochrome) to the phenol-red free media for 1h before imaging. For fixed cell samples, the cells were washed 1x in PBS, fixed for 15 min using 4% PFA in PBS, and washed 3x in PBS. For LatB-treated cells, 1 µg/ml latrunculin B (Cayman Chemicals, 10010631) was added to the cell culture medium for 1h before imaging. Right before imaging the coverslip with cells was attached to the bottom of the four transparent polished walls of a sliced commercial glass cuvette (704-000-20-10, Hellma) using two-part silicone rubber (Ecoflex 5, Reynolds Advanced Materials) as described previously^32^. All cell samples were imaged at 37°C and at 5% CO_2_ in fresh cell imaging media containing 330 nM SiR-DNA and 1 µg/ml latrunculin B in the case of LatB-treated cells.

### Optical setup

The optical setup was built around a conventional inverted microscope (IX71, Olympus) as described previously^32,42^ (Figure 1B). The illumination lasers (488 nm, 150 mW, CW, Coherent; 560 nm, 1000 mW, CW, MPB Communications; 647 nm, 120 mW, CW, Coherent) were spectrally filtered (488 nm: FF01-475/23-25 bandpass excitation filter; 561 nm: ff01-554/23-25 excitation filter; 647 nm: ff01-631/36-25 excitation filter, all Semrock), circularly polarized (LPVISB050-MP2 polarizers, Thorlabs; and 488 nm: Z-10-A-.250-B-488 quarter-wave plate, Tower Optical; 561 nm: WPQ05M-561 quarter-wave plate, Thorlabs; 647 nm: WPQ05M-633 quarter-wave plate, Thorlabs), and expanded and collimated using lens telescopes. Illumination was controlled with a shutter (VS14S2T1 with VMM-D3 three-channel driver, Vincent Associates Uniblitz) and synchronized with the detection optics via MATLAB. The lasers were either introduced into the epi-illumination pathway through a Köhler lens or sent to the light sheet illumination pathway; the pathway was easily switched with a flip mirror. The light sheet illumination pathway consisted of a cylindrical lens (ACY254-200-A, *f* = 200 mm, Thorlabs), which focused the light in only one dimension onto a motorized mirror (8821 mirror mount with 8742 Picomotor controller, Newport). The motorized mirror plane was imaged onto the back aperture of a long working distance illumination objective (378-803-3, 10x, NA 0.28, Mitutoyo) by two lenses in a 4*f* configuration. The illumination objective then focused the light sheet, which was directed into the sample chamber at an angle of about 10° using a mirror mounted on a custom-made holder attached to the illumination objective (Supplemental Figure S3). The entire light sheet illumination path was mounted on a breadboard above the microscope stage and could be moved by an xyz translation stage (460P, Newport).

The sample was mounted on an xy coarse translation stage (M26821LOJ, Physik Instrumente) and a precise xyz piezoelectric stage (P-545.3C7, Physik Instrumente). The light emitted from the fluorophores was detected by a high NA detection objective (UPLSAPO100XO, 100x, NA 1.4, Olympus), spectrally filtered (Di01-R405/488/561/635 dichroic, ZET488NF notch filter, ZET561NF notch filter, and ZET647NF notch filter, all Chroma, and green channel: FF03-525/50-25 bandpass filter, Semrock; red channel: ET700/75m bandpass filter, Chroma), and focused by the microscope tube lens, before entering a 4*f* imaging system. The first lens of the 4*f* system (*f* = 90 mm, G322389000, Qioptiq) was positioned one focal length from the intermediate image plane formed by the microscope tube lens. To create a two-channel 4f system a dichroic mirror (T660lpxrxt, Chroma) was inserted after the first 4f lens to reflect light with wavelengths shorter than 660 nm into one light path (“green channel”) and to transmit red light into a second light path (“red channel”). In the positions one focal length after the first 4*f* lens (i.e., the Fourier planes of the imaging paths), the phase of the emitted light was modulated to reshape the point spread function (PSF) to encode the axial position of the emitters using a transmissive, dielectric double-helix (DH) phase mask (534 nm, 2.7 mm diameter, Double-Helix Optics, LLC) in the green channel and a Tetrapod phase mask with 6 µm axial range (fabricated using standard photolithography methods at the Stanford Nanofabrication Facility as described in^32^) in the red channel. After phase modulation, the light was focused by second 4*f* lenses (*f* = 90 mm, G322389000, Qioptiq) and imaged using two EM-CCD cameras (green channel: iXon Ultra 897; red channel: iXon3 897, both Andor).

The entire microscope body and the light sheet breadboard were enclosed in a custom-built cage incubator in which the temperature and CO_2_ concentration were controlled (H201-T-UNIT-BL temperature control unit and CO2-UNIT-BL-ES CO_2_ controller with external sensor, both OkoLab). In all cell measurements the environment was set to 37°C and 5% CO_2_.

### Cell imaging

Images were acquired using either 20 ms exposure time and continuous acquisition (50 Hz) or 50 ms exposure time and time-lapsed acquisition with 2 s image intervals (0.5 Hz). Custom scripts were written in MATLAB to synchronize the laser shutters and the image acquisition on the cameras. Chromosomal loci were excited by 488 nm light sheet illumination at ∼100 W/cm^2^, fiducial beads were excited by 560 nm epiillumination at 5 W/cm^2^, and stained DNA was excited by 647 nm epi-illumination at 500 mW/cm^2^. All three lasers were on simultaneously for all frames. Dark frames were acquired when the lasers were off and the camera shutters were closed for both sets of imaging settings.

### Data analysis

A schematic outlining the main data analysis steps is shown in Supplemental Figure S1D. In brief, calibration scans were acquired of fluorescent beads using the DH PSF in the green channel and the Tetrapod PSF in the red channel. Registration images were acquired simultaneously in both color channels using clear aperture PSFs. The DH calibration scans were then used to localize the chromosomal loci in the green channel. The Tetrapod PSF calibrations were used to localize a fiducial bead at the coverslip in the red channel during cell imaging. The chromatin in the nucleus was imaged together with the fiducial bead in the red channel. For each frame acquired of the loci in the green channel, a frame was acquired in the red channel of the position of the fiducial bead and of the cell nucleus. The registration images were used to register the two color channels and the red channel data were then transformed into green coordinates. Finally, the transformed tracks of the fiducial bead and the cell nucleus were used to correct the chromosomal loci motion for drift and nuclear translation, respectively. Details of the different steps are described below.

### Tracking of chromosomal loci using the DH PSF

The labeled chromosomal loci were imaged using the DH PSF and localized using the software Easy-DHPSF v.2^54^. This software fits the lobes of each PSF using non-linear least-squares functions in MATLAB with a pair of identical, radially symmetric 2D Gaussians as the objective function. The mid-point between the two lobes encodes the lateral (xy) position of the locus and the angle between the lobes encodes the axial (z) position.

A calibration z-stack was acquired by imaging of 100 nm fluorescent beads (Tetraspeck, T7279, Invitrogen) at the coverslip surface while axially scanning over a 2-um range using the piezo stage. The beads were attached to the coverslip as described in the *Sample preparation* section and submerged in water to mimic cell imaging conditions. Dark frames were acquired with the same imaging settings while the shutter of the camera was closed. The z-stack and the dark frames were used to calibrate the DH PSF. The images of the chromosomal loci were then used to find suitable thresholds for locus fitting and the positions of each locus were then fitted in each camera frame. The localizations were filtered to have a distance between lobes between 4.1 px and 6.2 px (1 px is 160 nm) and to have a localization precision less than 30 nm in x, y, and z. All z localizations were rescaled by a factor of 0.75 to account for the refractive index mismatch between glass and the sample^55^.

The individual localizations were linked into time trajectories using custom software based on a nearest-neighbor-type particle tracking heuristic with filters for maximum distance moved between frames, maximum number of off frames in which the locus was not detected, and minimum number of total frames per track. Each track was inspected by eye to ensure correct linking of localizations.

### Tracking of sample drift using fiducial beads and the Tetrapod PSF

In order to calibrate the experimental Tetrapod PSF, an axial scan of a 100 nm fluorescent bead (Tetraspeck, T7279, Invitrogen) at the coverslip surface was performed over a 7 µm range using the piezo stage. The bead was attached to the coverslip as described in the *Sample preparation* section and submerged in water to mimic cell imaging conditions. A model of the Tetrapod based on vectorial diffraction theory was modified with a phase term in the Fourier plane consisting of a linear combination of Zernike polynomials (first six orders). The contributions of the Zernike modes were optimized using a maximum likelihood-based phase retrieval procedure described previously^56,57^ in order to produce a modified PSF model tailored to the experimental Tetrapod measurements.

Fiducial beads at or near the coverslip (z ≈ 0) in cell samples were tracked in 3D (x, y, and nominal focal plane position f) by manual detection of the region of interest (ROI). A library of Tetrapod PSF images was simulated at 50 nm intervals in nominal focal position. In the first frame of each fiducial trajectory, the coarse axial position was estimated by normalizing the ROI and convolving it with each normalized, simulated image using the *conv2* function in MATLAB. Fine localization was performed using maximum-likelihood estimation by comparing the frame to a 3D spline interpolation of the simulated PSF stack. A Poisson noise model was used for the maximum likelihood objective function. Outputs from the fine localization of frame n were supplied as inputs to frame n+1, and in the case of large lateral drift the ROI location within the field of view was updated.

### Tracking of nuclear translation using nuclear center of mass

Live cells migrate on the coverslip and the translational motion of the cell nucleus should be subtracted to properly report the motion of the loci within the nucleus. To measure the nuclear translation, an image was acquired of the entire stained nucleus in the red channel for each image of chromosomal loci acquired in the green channel. The images of the nucleus were then analyzed using Fiji. All images were converted into binary images, the center of mass of each binary image was measured, and the resulting center of mass coordinates were imported into MATLAB.

### Correcting the chromosomal locus tracks for drift and nuclear translation

All chromosomal loci tracks were corrected for 3D sample drift using the measured motion of fiducial beads (Supplemental Figure S1). For long-term tracking (0.5 Hz), the tracks were also corrected in x and y for any nuclear translation caused by for example cell migration. For tracking at faster time scales (50 Hz), the nuclear translation was negligible for our analysis. We therefore chose to use the fiducial beads to correct for drift in x, y, and z for these data, since the fiducial bead had better localization precision. Additional correction for nuclear translation would here risk adding more noise than correction value.

The fiducial beads and the nuclei were imaged in the red channel, while the chromosomal loci were tracked in the green channel. Before correction, the red and green channels thus had to be registered and the data from the red channel transformed into the green coordinates. Registration images were acquired of 100 nm fluorescent bead (Tetraspeck, T7279, Invitrogen) at the coverslip surface simultaneously in both color channels using clear aperture PSFs. The registration images were imported into Fiji, where the background was subtracted using the rolling ball method. The red channel image was flipped horizontally to facilitate image registration. The image from the red channel was then registered onto the green channel image in MATLAB using an affine transform with the function *imregtform*, and the resulting transformation matrix was saved.

The resulting transformation matrix was then used to transform the fiducial and nuclear data from the red to the green channel coordinates using the *transformPointsForward* function in MATLAB. The transformed drift and nuclear data were then smoothed in MATLAB using the *rlowess* method over a window of 20 data points to reduce the impact of noise before subtraction from the corresponding chromosomal loci data.

### MSD analysis

The most ubiquitous statistical measure used to analyze single-particle tracking data, including the dynamics of chromosomal loci, is the MSD. In this work we used an expression for fitting of the experimental MSD curves that accounts for both static errors (due to finite localization precision) and dynamic errors (motion blur due to finite exposure time) developed for anomalous diffusion consistent with fractional Brownian motion^43,58^:

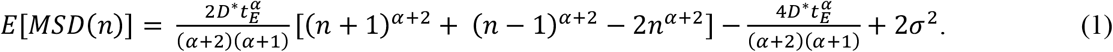

Here *E* refers to the expectation value of MSD(n), t_E_ is the exposure time, n is the number of frames spanning a lag, D* is the effective diffusion coefficient, α∈(0,2], and 2*σ*^2^ is the static error term also including the spreading of the PSF due to movement of the locus. The second to last term describes the effects of the dynamic error.

We fitted the first 15 lags of both the ensemble time-averaged MSDs and the single-track time-averaged MSDs to this equation in MATLAB with free parameters D*, α, and *σ*. Fits were performed by minimizing the square error in log-space^43^. The choice of 15 lags (300 ms) resulted in a reasonable tradeoff between having enough data points for robust fitting and keeping the time scale of the analysis short. Since each track consisted of many hundreds of localizations, we were always well below the typical criterion of only analyzing time lags less than ¼ of the total number of localizations^59^. We imposed physically reasonable yet modest lower and upper bounds, where the parameters were constrained: D* to (0, 1) µm^2^/s^α^, α to (0, 2), and *σ* to (0.001, 100) nm. Tracks resulting in α < 0.05 or α > 1.95 typically represented bad fits and such tracks were excluded from the analysis.

### Validation of methodology by tracking of beads in a glycerol-water mixture

As a control of the methodology, 100 nm fluorescent beads (505/515 nm, F8803, Invitrogen) in a glycerol-water mixture were tracked using the same apparatus and analyzed using the same methodology as the cell samples (Supplemental Figure S6). All z localizations were rescaled by a factor of 0.957 to account for the refractive index mis-match between glass and glycerol mixture^60^. In these experiments the fiducial beads at the coverslip were used for drift correction in x, y, and z. A total of 24 beads were tracked and each track consisted of at least 1300 frames. All data was acquired at 50 Hz.

### Validation of results from simulations

In order to demonstrate the validity of our analysis, as well as to benchmark the robustness of the analysis to track parameters and initial conditions, we simulated tracks of fractional Brownian motion using the method of circulant embedding^43,61^. The simulated tracks were sampled with a constant time interval which is the exposure time divided by the number of photons per frame. Then, a Gaussian noise term representing the localization precision was added to each true position and the centroid of photon positions was used as the estimated molecule position. For our analysis, we simulated 50 tracks independently in each of the three dimensions using conditions similar to the experimental conditions: 20 ms exposure time, 50,000 photons per frame, *σ* = 15 nm, track lengths of 650 or 1000 frames, D*_sim_ = 1×10^−3^ µm^2^/s^α^, and α _sim_ = 0.5.

### Statistical analysis

To compare the distributions shown in Figure 4, p-values were calculated based on a Mann-Whitney U test, which is a nonparametric statistical significance test.

### Code availability

The custom-written code generated during the current study is available from the corresponding author upon request. Calibration and fitting of Tetrapod fiducial images for drift correction was performed using a modified version of the open-source Easy-Pupil-Finder software^56^ (https://sourceforge.net/projects/easy-pupil-finder/). Calibration and fitting analysis of DH-PSF images was performed using a modified version of the open-source Easy-DHPSF v.2 software^54^ (https://sourceforge.net/projects/easy-dhpsf/).

### Data availability

The data generated and analyzed during the current study are available from the corresponding author upon request.

## ACKNOWLEDGMENTS

The authors thank Peter D. Dahlberg and Annina Sartor for their help constructing the cage incubator and fabricating the light sheet mirror holder. The authors thank Camille A. Bayas for helpful discussions regarding the data analysis. This work was supported in part by the National Institute of General Medical Sciences (Grant No. R35GM118067) to W.E.M. and by the National Institute of Biomedical Imaging and Bioengineering (Grant No. U01EB021237) to W.E.M. A.-K.G. acknowledges partial financial support from National Institute of General Medical Sciences of the National Institutes of Health (Grant No. R00GM134187), the Welch Foundation (Grant No. C-2064-20210327), and startup funds from the Cancer Prevention and Research Institute of Texas (Grant No. RR200025). P.N.P. acknowledges support from a Bio-X Stanford Interdisciplinary Graduate Fellowship.

## AUTHOR CONTRIBUTIONS

A-.K.G and P.N.P designed the experimental setup. A-.K.G. and R.P.G designed the experiments and wrote the manuscript. R.P.G. generated the array and guide plasmids and the cell line. A-.K.G. performed the experiments and analyzed all data except the Tetrapod data. P.N.P. analyzed the Tetrapod data. J.T.L. and W.E.M. supervised the work. All authors discussed the data and edited the manuscript.

## REFERENCES

1. Cremer, T. & Cremer, C. Chromosome territories, nuclear architecture and gene regulation in mammalian cells. Nat Rev Genet 2, 292–301 (2001).

2. Lieberman-Aiden, E. et al. Comprehensive Mapping of Long-Range Interactions Reveals Folding Principles of the Human Genome. Science 326, 289 (2009).

3. Parmar, J. J., Woringer, M. & Zimmer, C. How the Genome Folds: The Biophysics of Four-Dimensional Chromatin Organization. Annu. Rev. Biophys. 48, 231–253 (2019).

4. Chen, B. et al. Dynamic Imaging of Genomic Loci in Living Human Cells by an Optimized CRISPR/Cas System. Cell 155, 1479–1491 (2013).

5. Tanenbaum, M. E., Gilbert, L. A., Qi, L. S., Weissman, J. S. & Vale, R. D. A Protein-Tagging System for Signal Amplification in Gene Expression and Fluorescence Imaging. Cell 159, 635–646 (2014).

6. Kamiyama, D. et al. Versatile protein tagging in cells with split fluorescent protein. Nat Commun 7, 11046 (2016).

7. Bertrand, E. et al. Localization of ASH1 mRNA Particles in Living Yeast. Molecular Cell 2, 437–445 (1998).

8. Hocine, S., Raymond, P., Zenklusen, D., Chao, J. A. & Singer, R. H. Single-molecule analysis of gene expression using two-color RNA labeling in live yeast. Nat Methods 10, 119–121 (2013).

9. Viswanathan, S. et al. High-performance probes for light and electron microscopy. Nat Methods 12, 568–576 (2015).

10. Kirchhofer, A. et al. Modulation of protein properties in living cells using nanobodies. Nat Struct Mol Biol 17, 133–138 (2010).

11. Ghosh, R. P. et al. A fluorogenic array for temporally unlimited single-molecule tracking. Nat. Chem. Biol. 15, 401–409 (2019).

12. Levi, V., Ruan, Q., Plutz, M., Belmont, A. S. & Gratton, E. Chromatin Dynamics in Interphase Cells Revealed by Tracking in a Two-Photon Excitation Microscope. Biophysical Journal 89, 4275–4285 (2005).

13. Katayama, Y. et al. Real-time nanomicroscopy via three-dimensional single-particle tracking. ChemPhysChem 10, 2458–2464 (2009).

14. Perillo, E. P. et al. Deep and high-resolution three-dimensional tracking of single particles using nonlinear and multiplexed illumination. Nat. Commun. 6, 7874 (2015).

15. Toprak, E., Balci, H., Blehm, B. H. & Selvin, P. R. Three-Dimensional Particle Tracking via Bifocal Imaging. Nano Lett. 7, 2043–2045 (2007).

16. Ram, S., Prabhat, P., Chao, J., Sally Ward, E. & Ober, R. J. High accuracy 3D quantum dot tracking with multifocal plane microscopy for the study of fast intracellular dynamics in live cells. Biophys. J. 95, 6025–6043 (2008).

17. Juette, M. F. et al. Three-dimensional sub–100 nm resolution fluorescence microscopy of thick samples. Nat. Methods 5, 527–529 (2008).

18. Ram, S., Kim, D., Ober, R. J. & Ward, E. S. 3D single molecule tracking with multifocal plane microscopy reveals rapid intercellular transferrin transport at epithelial cell barriers. Biophys. J. 103, 1594–1603 (2012).

19. Abrahamsson, S. et al. Fast multicolor 3D imaging using aberration-corrected multifocus microscopy. Nat. Methods 10, 60–63 (2013).

20. Chen, J. et al. Single-molecule dynamics of enhanceosome assembly in embryonic stem cells. Cell 156, 1274–1285 (2014).

21. Knight, S. C. et al. Dynamics of CRISPR-Cas9 genome interrogation in living cells. Science 350, 823–826 (2015).

22. Smith, C. S. et al. Nuclear accessibility of β-actin mRNA is measured by 3D single-molecule real-time tracking. Journal of Cell Biology 209, 609–619 (2015).

23. von Diezmann, L., Shechtman, Y. & Moerner, W. E. Three-dimensional localization of single molecules for super-resolution imaging and single-particle tracking. Chem. Rev. 117, 7244–7275 (2017).

24. Kao, H. P. & Verkman, A. S. Tracking of single fluorescent particles in three dimensions: use of cylindrical optics to encode particle position. Biophys. J. 67, 1291–1300 (1994).

25. Huang, B., Jones, S. A., Brandenburg, B. & Zhuang, X. Whole-cell 3D STORM reveals interactions between cellular structures with nanometer-scale resolution. Nat. Methods 5, 1047–1052 (2008).

26. Spille, J.-H., Kaminski, T., Königshoven, H.-P. & Kubitscheck, U. Dynamic three-dimensional tracking of single fluorescent nanoparticles deep inside living tissue. Opt. Express 20, 19697 (2012).

27. Li, Y., Hu, Y. & Cang, H. Light sheet microscopy for tracking single molecules on the apical surface of living cells. J. Phys. Chem. B 117, 15503–15511 (2013).

28. Izeddin, I. et al. Single-molecule tracking in live cells reveals distinct target-search strategies of transcription factors in the nucleus. eLife 3, e02230 (2014).

29. Backer, A. S., Backlund, M. P., von Diezmann, A. R., Sahl, S. J. & Moerner, W. E. A bisected pupil for studying single-molecule orientational dynamics and its application to three-dimensional super-resolution microscopy. Appl. Phys. Lett. 104, 193701 (2014).

30. Jia, S., Vaughan, J. C. & Zhuang, X. Isotropic three-dimensional super-resolution imaging with a self-bending point spread function. Nat. Photonics 8, 302–306 (2014).

31. Lew, M. D., Lee, S. F., Badieirostami, M. & Moerner, W. E. Corkscrew point spread function for far-field three-dimensional nanoscale localization of pointlike objects. Opt. Lett. 36, 202–204 (2011).

32. Gustavsson, A.-K., Petrov, P. N., Lee, M. Y., Shechtman, Y. & Moerner, W. E. 3D single-molecule super-resolution microscopy with a tilted light sheet. Nat. Commun. 9, 123 (2018).

33. Pavani, S. R. P. et al. Three-dimensional, single-molecule fluorescence imaging beyond the diffraction limit by using a double-helix point spread function. Proc. Natl. Acad. Sci. U.S.A. 106, 2995–2999 (2009).

34. Thompson, M. A., Casolari, J. M., Badieirostami, M., Brown, P. O. & Moerner, W. E. Three-dimensional tracking of single mRNA particles in Saccharomyces cerevisiae using a double-helix point spread function. Proceedings of the National Academy of Sciences 107, 17864–17871 (2010).

35. Backlund, M. P., Joyner, R., Weis, K. & Moerner, W. E. Correlations of three-dimensional motion of chromosomal loci in yeast revealed by the double-helix point spread function microscope. Mol. Biol. Cell 25, 3619–3629 (2014).

36. Shechtman, Y., Sahl, S. J., Backer, A. S. & Moerner, W. E. Optimal point spread function design for 3D imaging. Phys. Rev. Lett. 113, 133902 (2014).

37. Shechtman, Y., Weiss, L. E., Backer, A. S., Sahl, S. J. & Moerner, W. E. Precise three-dimensional scan-free multiple-particle tracking over large axial ranges with tetrapod point spread functions. Nano Lett. 15, 4194–4199 (2015).

38. Shechtman, Y. et al. Observation of live chromatin dynamics in cells via 3D localization microscopy using Tetrapod point spread functions. Biomed. Opt. Express 8, 5735 (2017).

39. Weiss, L. E. et al. Three-dimensional localization microscopy in live flowing cells. Nat. Nanotechnol. 15, 500–506 (2020).

40. Huisken, J., Swoger, J., Del Bene, F., Wittbrodt, J. & Stelzer, E. H. K. Optical sectioning deep inside live embryos by selective plane illumination microscopy. Science 305, 1007 (2004).

41. Hu, Y. S., Zimmerley, M., Li, Y., Watters, R. & Cang, H. Single-molecule super-resolution light-sheet microscopy. ChemPhysChem 15, 577–586 (2014).

42. Gustavsson, A.-K., Petrov, P. N. & Moerner, W. E. Light sheet approaches for improved precision in 3D localization-based super-resolution imaging in mammalian cells [Invited]. Opt. Express 26, 13122 (2018).

43. Backlund, M. P., Joyner, R. & Moerner, W. E. Chromosomal locus tracking with proper accounting of static and dynamic errors. Phys. Rev. E 91, 062716 (2015).

44. Petrov, P. N. & Moerner, W. E. Addressing systematic errors in axial distance measurements in single-emitter localization microscopy. Opt. Express 28, 18616 (2020).

45. Molenaar, C. Visualizing telomere dynamics in living mammalian cells using PNA probes. The EMBO Journal 22, 6631–6641 (2003).

46. Jegou, T. et al. Dynamics of Telomeres and Promyelocytic Leukemia Nuclear Bodies in a Telomerase-negative Human Cell Line. MBoC 20, 2070–2082 (2009).

47. Bronstein, I. et al. Transient Anomalous Diffusion of Telomeres in the Nucleus of Mammalian Cells. Phys. Rev. Lett. 103, 018102 (2009).

48. Bronshtein, I. et al. Loss of lamin A function increases chromatin dynamics in the nuclear interior. Nat. Commun 6, 8044 (2015).

49. Kepten, E., Weron, A., Bronstein, I., Burnecki, K. & Garini, Y. Uniform Contraction-Expansion Description of Relative Centromere and Telomere Motion. Biophysical Journal 109, 1454–1462 (2015).

50. Shinkai, S., Nozaki, T., Maeshima, K. & Togashi, Y. Dynamic Nucleosome Movement Provides Structural Information of Topological Chromatin Domains in Living Human Cells. PLoS Comput Biol 12, e1005136 (2016).

51. Bronshtein, I. et al. Exploring chromatin organization mechanisms through its dynamic properties. Nucleus 7, 27–33 (2016).

52. Qin, P. et al. Live cell imaging of low-and non-repetitive chromosome loci using CRISPR-Cas9. Nat Commun 8, 14725 (2017).

53. Makhija, E., Jokhun, D. S. & Shivashankar, G. V. Nuclear deformability and telomere dynamics are regulated by cell geometric constraints. Proc Natl Acad Sci USA 113, E32–E40 (2016).

54. Bayas, C. A., Diezmann, A. von, Gustavsson, A.-K. & Moerner, W. E. Easy-DHPSF 2.0: open-source software for three-dimensional localization and two-color registration of single molecules with nanoscale accuracy. Protocol Exchange (2019) doi:10.21203/rs.2.9151/v2.

55. Li, Y. et al. Real-time 3D single-molecule localization using experimental point spread functions. Nat Methods 15, 367–369 (2018).

56. Petrov, P. N., Shechtman, Y. & Moerner, W. E. Measurement-based estimation of global pupil functions in 3D localization microscopy. Opt. Express 25, 7945–7959 (2017).

57. Siemons, M., Hulleman, C. N., Thorsen, R. Ø., Smith, C. S. & Stallinga, S. High precision wavefront control in point spread function engineering for single emitter localization. http://biorxiv.org/lookup/doi/10.1101/267864 (2018) doi:10.1101/267864.

58. Savin, T. & Doyle, P. S. Static and Dynamic Errors in Particle Tracking Microrheology. Biophysical Journal 88, 623–638 (2005).

59. Saxton, M. J. & Jacobson, K. Single-particle tracking: Applications to Membrane Dynamics. Annu. Rev. Biophys. Biomol. Struct. 26, 373–399 (1997).

60. Takamura, K., Fischer, H. & Morrow, N. R. Physical properties of aqueous glycerol solutions. Journal of Petroleum Science and Engineering 98–99, 50–60 (2012).

61. Stochastic Geometry, Spatial Statistics and Random Fields: Models and Algorithms. vol. 2120 (Springer International Publishing, 2015).

