## Supplemental Information for "Fast and parallel nanoscale 3D tracking of heterogeneous mammalian chromatin dynamics"

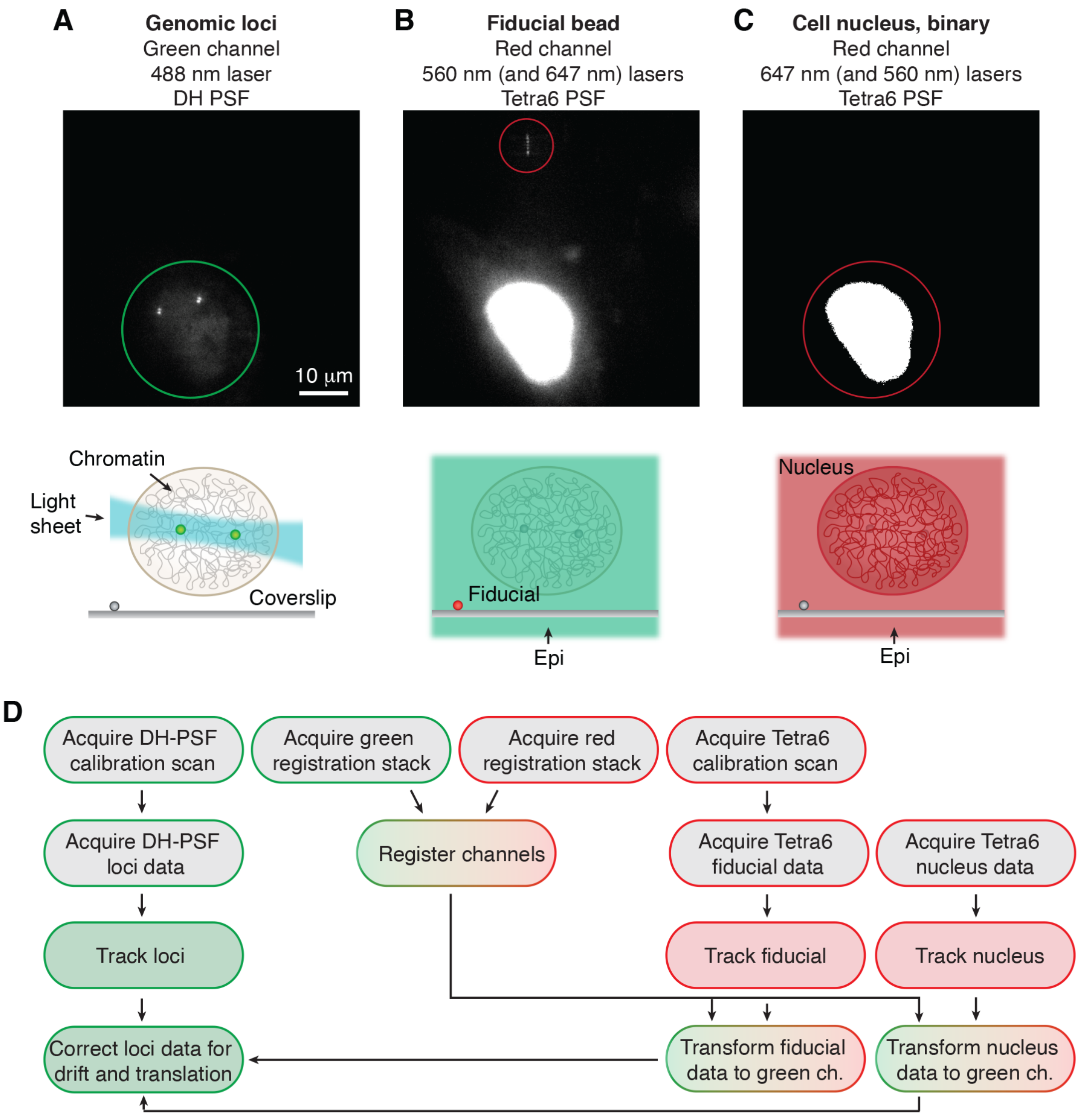


Supplemental Figure S1. Schematic showing the excitation, detection, and analysis schemes. (A) Labeled chromosomal loci were excited by 488 nm light sheet illumination and detected in the green channel using the double-helix (DH) PSF. (B) Fiducial beads melted to the coverslip were excited by 560 nm epi-illumination and detected in the red channel using the Tetrapod PSF with 6 μm axial range (Tetra6 PSF). The detected bead motion was used to correct for sample drift. (C) The SiR-DNA labeled chromatin was excited by 647 nm epi-illumination and detected in the red channel using the Tetra6 PSF. Detected nuclear motion was used to correct for nuclear translation. All lasers were turned on for each frame and one frame in the red channel was acquired for each frame acquired in the green channel. (D) Schematic showing the analysis workflow.

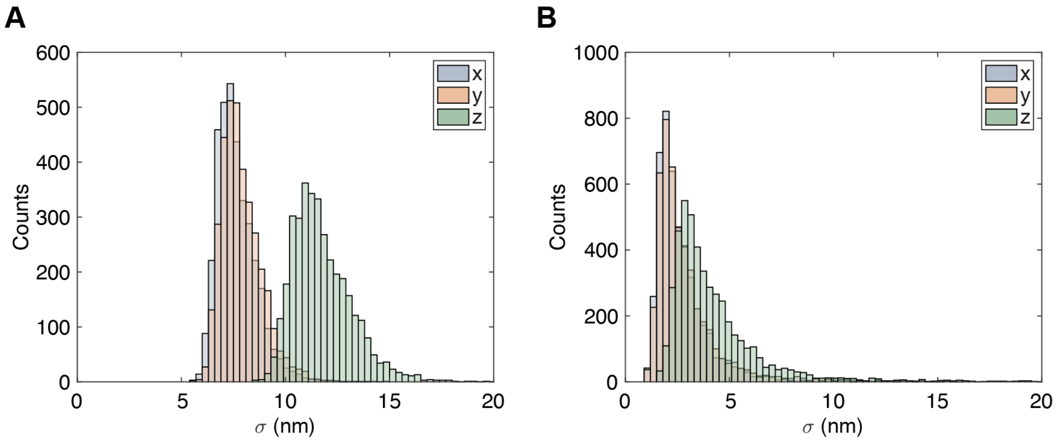


Supplemental Figure S2. Localization precision. Histograms showing the distribution of localization precisions in x, y, and z for localizations over 5000 frames when tracking a locus at (A) 50 Hz (20 ms exposure time) and (B) 0.5 Hz (50 ms exposure time with time-lapsing).

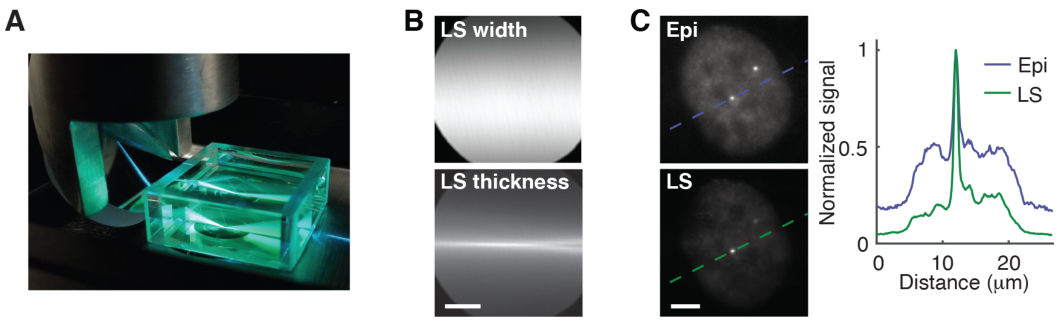


Supplemental Figure S3. Light sheet design and characterization. (A) Photo showing the light sheet being reflected by a mirror mounted on an aluminum tube attached to the illumination objective. The light sheet is introduced into the sample chamber at an angle and focused at the coverslip at the bottom of the chamber. (B) The light sheet (LS) width and thickness imaged when the light traveled through a fluorescent solution containing 20 μM Chromeo 488. The light sheet thickness was imaged by turning the cylindrical lens by 90°. Scale bar is 20 μm. By fitting the images to Gaussian functions, the width and thickness were determined to be 46 μm ($1/e^2$) and 2.5 μm ($1/e^2$), respectively. (C) Comparison between epi-illumination (Epi) and light sheet illumination (LS) for excitation of labeled chromosomal loci in a cell nucleus. The graph shows improvement in contrast by a factor of more than 2.5 of when using light sheet illumination. Scale bar is 5 μm.

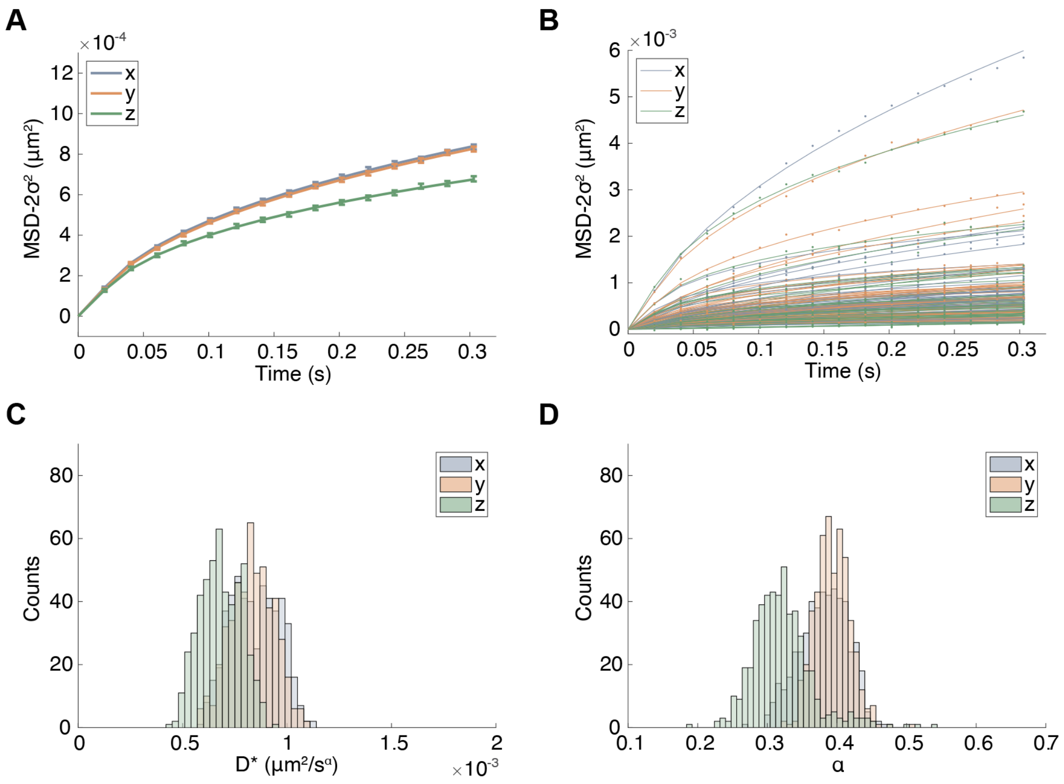


Supplemental Figure S4. MSD analysis of experimental tracks in untreated cells. (A) Experimental data and fits of the time-averaged ensemble MSDs in x, y, and z. The points show the mean experimental data with error bars showing the SEM from 100 bootstraps with 50% dropout. The complete data consisted of 54 tracks, each with a minimum track length of 640 frames. (B) Experimental data and fits of MSDs of the individual tracks in x, y, and z. (C) Histograms of the resulting D* from fits of the time-averaged ensemble MSDs using 500 bootstraps with 50% dropout. (D) Histograms of the resulting α from fits of the time-averaged ensemble MSDs using 500 bootstraps with 50% dropout. The resulting estimated parameters using 500 bootstraps with 50% dropout were $D^*_{x,y,z}$ = [0.85±0.01, 0.83±0.01, 0.68±0.01]×$10^{-3}$ $\mu m^2/s^{\alpha}$ and $\alpha_{x,y,z}$ = [0.379±0.003, 0.389±0.003, 0.312±0.004].

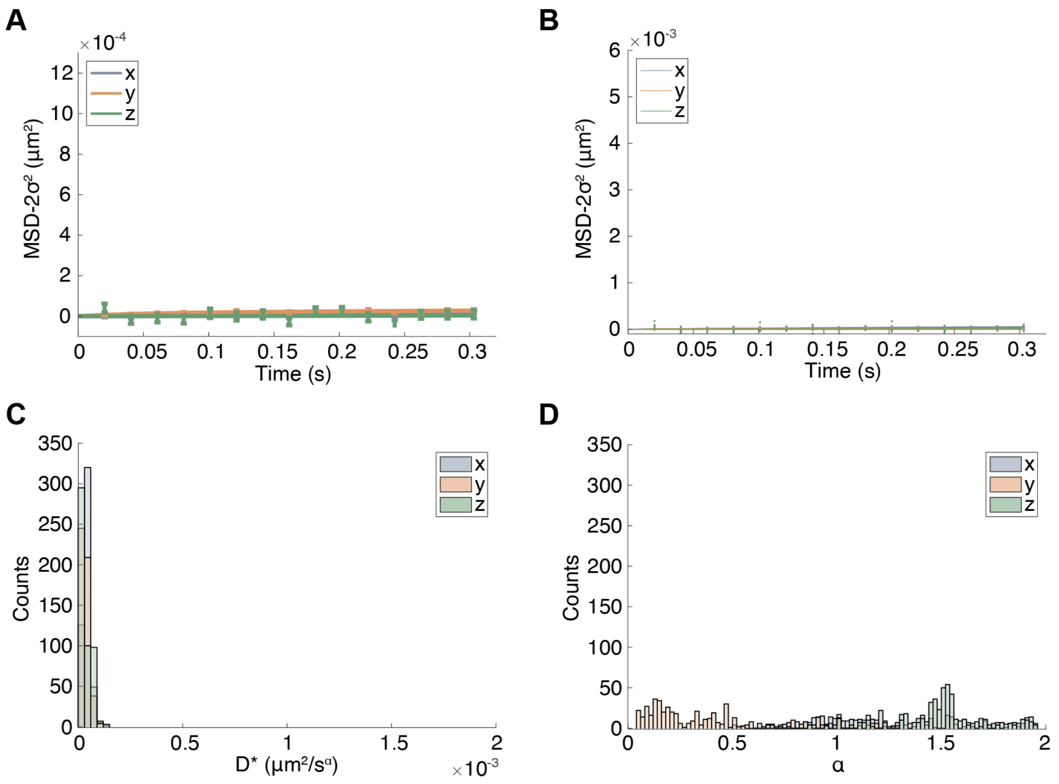


Supplemental Figure S5. MSD analysis of tracks in fixed cells. (A) Experimental data and fits of the time-averaged ensemble MSDs in x, y, and z. The points show the mean experimental data with error bars showing the SEM from 100 bootstraps with 50% dropout. The complete data consisted of 10 tracks, each with a minimum track length of 640 frames. (B) Experimental data and fits of MSDs of the individual tracks in x, y, and z. (C) Histograms of the resulting D* from fits of the time-averaged ensemble MSDs using 500 bootstraps with 50% dropout. (D) Histograms of the resulting α from fits of the time-averaged ensemble MSDs using 500 bootstraps with 50% dropout. The resulting estimated parameters using 500 bootstraps with 50% dropout were $D^*_{x,y,z}$ = [3.64±0.14, 3.21±0.19, 1.22±0.28]$\times 10^{-5}$ $\mu m^2/s^{\alpha}$ and $\alpha_{x,y,z}$ = [1.25±0.03, 0.19±0.03, 1.57 ±0.03].

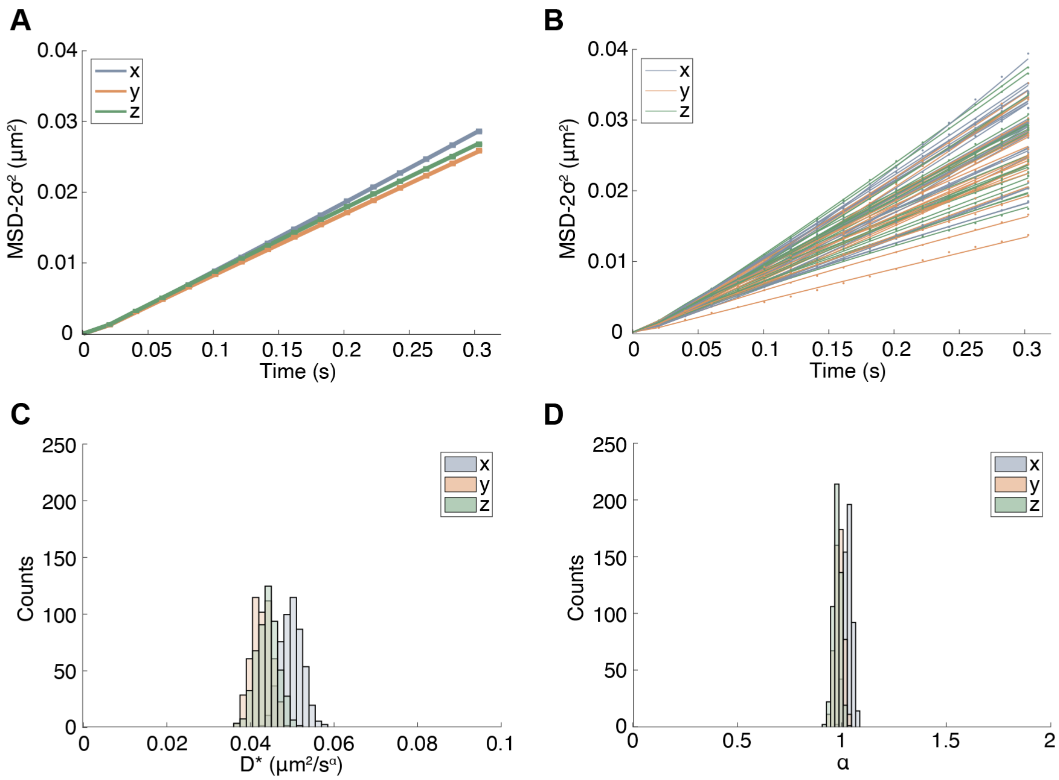


Supplemental Figure S6. MSD analysis of Brownian motion of beads diffusing in a glycerol mixture results in isotropic motion. (A) Fits of the time-averaged ensemble MSDs in x, y, and z. The points show the mean experimental data with error bars showing the SEM from 100 bootstraps with 50% dropout. The data consists of 24 beads each with a minimum track length of 1300 frames. (B) Experimental data and fits of MSDs of the individual tracks in x, y, and z. (C) Histograms of the resulting D* from fits of the time-averaged ensemble MSDs using 500 bootstraps with 50% dropout. (D) Histograms of the resulting α from fits of the time-averaged ensemble MSDs using 500 bootstraps with 50% dropout. All data was acquired at 50 Hz and is corrected for index-mismatch between glass and glycerol using a z-correction factor of 1.453/1.518. The data was fit using initial guesses for fitting of $D^*_{init} = 1\times10^{-2}$ μm²/s$^{\alpha}$ and $\alpha_{init} = 1.0$. The resulting estimated parameters using 500 bootstraps with 50% dropout were $D^*_{x,y,z}$ = [4.97±0.03, 4.27±0.02, 4.40±0.03]$\times10^{-2}$ $\mu$m²/s$^{\alpha}$ and $\alpha_{x,y,z}$ = [1.024±0.002, 0.982±0.002, 0.972±0.002]. The estimated ensemble average D*is in reasonable agreement with that predicted from the Stokes-Einstein relation for an 85% (w/w) glycerol mixture (~$4.4\times10^{-2}$ $\mu$m²/s).

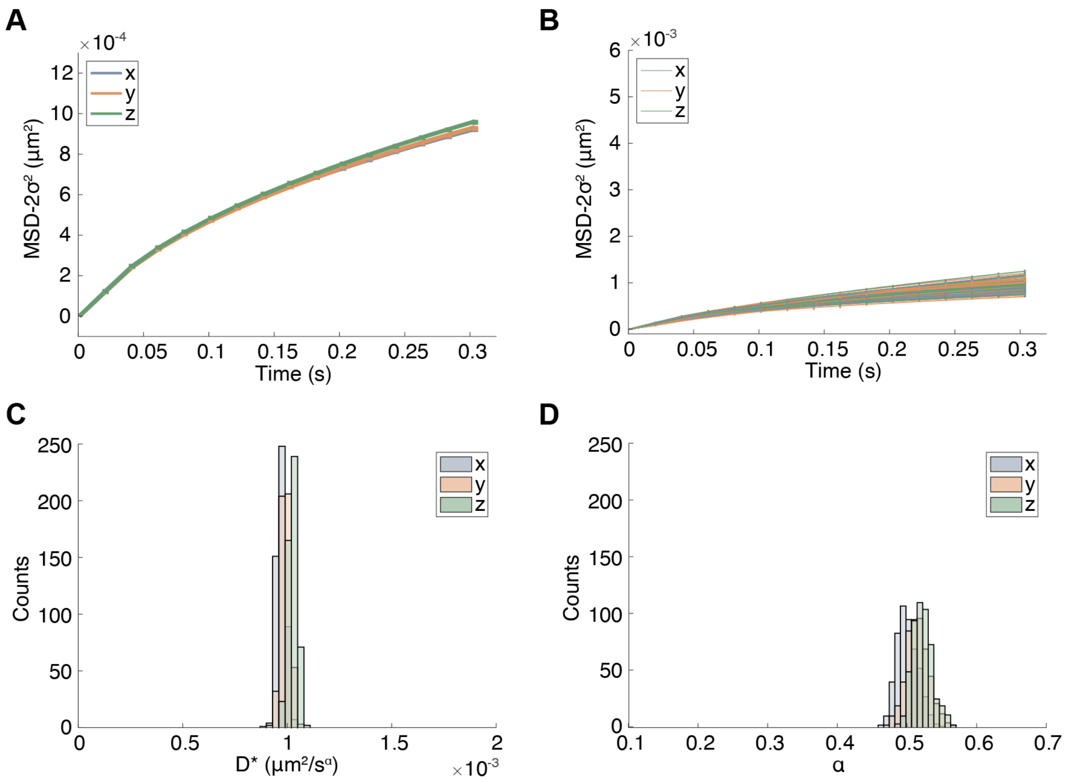


Supplemental Figure S7. Validation of analysis procedure by MSD analysis of simulated fractional Brownian motion using similar parameters for track generation and fitting as for the experimental data. 50 tracks each containing 1000 localizations were simulated using $D^*_{sim} = 1\times10^{-3}$ $\mu m^2/s^\alpha$ and $\alpha_{sim} = 0.5$, and fit using the same initial guesses as used for the experimental data: $D^*_{init} = 1\times10^{-3}$ $\mu m^2/s^\alpha$ and $\alpha_{init} = 0.5$. (A) Fits of the time-averaged ensemble MSDs, where E1-E3 represent the simulated motion in x, y, z, respectively. The points show the mean simulated data with error bars showing the SEM from 100 bootstraps with 50% dropout. (B) Experimental data and fits of MSDs of the individual tracks in x, y, and z. (C) Histograms of the resulting D* from fits of the time-averaged ensemble MSDs using 500 bootstraps with 50% dropout. (D) Histograms of the resulting α from fits of the time-averaged ensemble MSDs using 500 bootstraps with 50% dropout. The resulting estimated parameters using 500 bootstraps with 50% dropout were $D^*_{1,2,3}$ = [0.966 ± 0.002, 0.989 ± 0.002, 1.020 ± 0.002]$\times10^{-3}$ $\mu m^2/s^\alpha$ and $\alpha_{1,2,3}$ = [0.486 ± 0.002, 0.509 ± 0.002, 0.516 ± 0.002]. There is some heterogeneity between individual tracks, as expected from randomly simulated tracks with finite track length. However, the estimated ensemble average parameters are in good agreement with the simulated values.

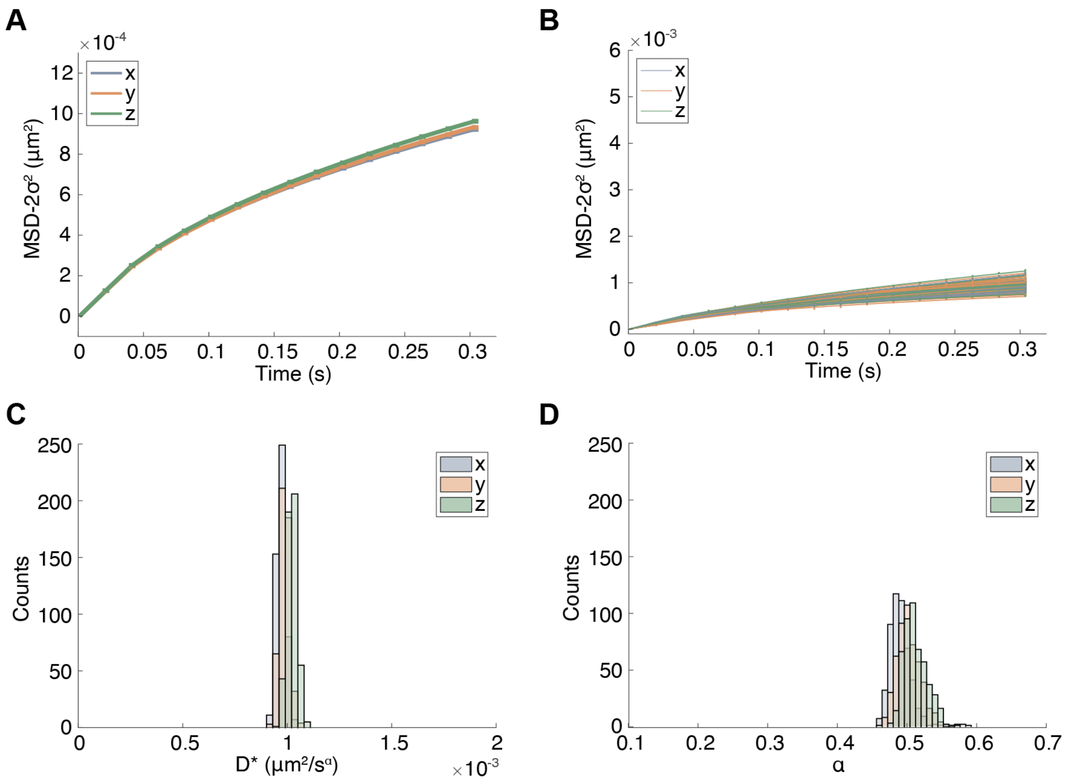


Supplemental Figure S8. The effect of varying the initial guesses of the fits. MSD analysis of simulated fractional Brownian motion using similar parameters for track generation as for the experimental data but using different initial guesses for the fits. 50 tracks each containing 1000 localizations were simulated using $D^*_{sim} = 1\times10^{-3}$ $\mu m^2/s^\alpha$ and $\alpha_{sim} = 0.5$, and fit using the initial guesses: $D^*_{init} = 2\times10^{-3}$ $\mu m^2/s^\alpha$ and $\alpha_{init} = 1.0$. (A) Fits of the time-averaged ensemble MSDs, where E1-E3 represent the simulated motion in x, y, z, respectively. The points show the mean simulated data with error bars showing the SEM from 100 bootstraps with 50% dropout. (B) Experimental data and fits of MSDs of the individual tracks in x, y, and z. (C) Histograms of the resulting D* from fits of the time-averaged ensemble MSDs using 500 bootstraps with 50% dropout. (D) Histograms of the resulting α from fits of the time-averaged ensemble MSDs using 500 bootstraps with 50% dropout. The resulting estimated parameters using 500 bootstraps with 50% dropout were $D^*_{1,2,3} = [0.964 \pm 0.002, 0.983 \pm 0.002, 1.020 \pm 0.002]\times10^{-3}$ $\mu m^2/s^\alpha$ and $\alpha_{1,2,3} = [0.482 \pm 0.001, 0.496 \pm 0.001, 0.502 \pm 0.002]$. Our analysis showed that the resulting parameters are robust with regards to the choice of initial guesses.

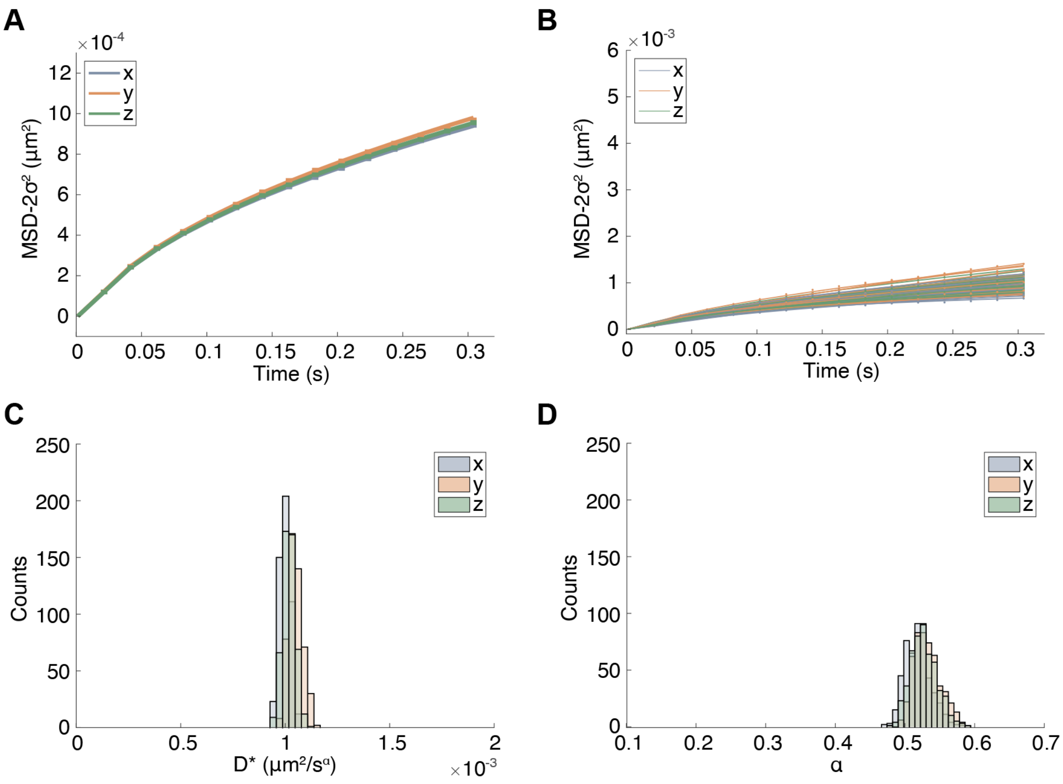


Supplemental Figure S9. The effect of having fewer localizations per track. MSD analysis of simulated fractional Brownian motion using similar parameters for track generation and fitting as the experimental data but using only 650 localizations per track. 50 tracks each containing 650 localizations were simulated using $D^*_{sim} = 1\times10^{-3}$ $\mu m^2/s^\alpha$ and $\alpha_{sim} = 0.5$, and fit using the same initial guesses as used for the experimental data: $D^*_{init} = 1\times10^{-3}$ $\mu m^2/s^\alpha$ and $\alpha_{init} = 0.5$. (A) Fits of the time-averaged ensemble MSDs, where E1-E3 represent the simulated motion in x, y, z, respectively. The points show the mean simulated data with error bars showing the SEM from 100 bootstraps with 50% dropout. (B) Experimental data and fits of MSDs of the individual tracks in x, y, and z. (C) Histograms of the resulting D* from fits of the time-averaged ensemble MSDs using 500 bootstraps with 50% dropout. (D) Histograms of the resulting α from fits of the time-averaged ensemble MSDs using 500 bootstraps with 50% dropout. The resulting estimated parameters using 500 bootstraps with 50% dropout were $D^*_{1,2,3}$ = [0.999 ± 0.003, 1.048 ± 0.003, 1.020 ± 0.003]$\times10^{-3}$ $\mu m^2/s^\alpha$ and $\alpha_{1,2,3}$ = [0.515 ± 0.002, 0.525 ± 0.002, 0.523 ± 0.002]. A reduction of the number of localizations per track leads, as expected, to a wider distribution in the estimated parameters, but the ensemble average values are still reasonably robust and in good agreement with the simulated values.

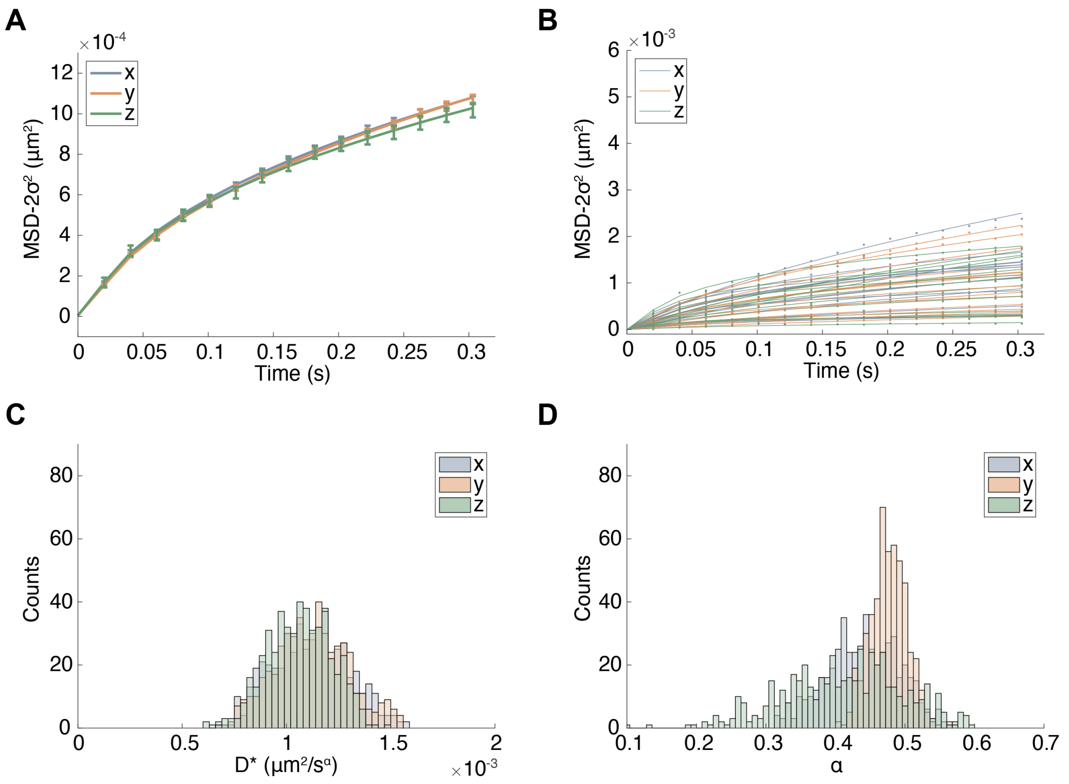


Supplemental Figure S10. MSD analysis of tracks in cells treated with Latrunculin B. (A) Experimental data and fits of the time-averaged ensemble MSDs in x, y, and z. The points show the mean experimental data with error bars showing the SEM from 100 bootstraps with 50% dropout. The complete data consisted of 14 tracks, each with a minimum track length of 640 frames. (B) Experimental data and fits of MSDs of the individual tracks in x, y, and z. (C) Histograms of the resulting D* from fits of the time-averaged ensemble MSDs using 500 bootstraps with 50% dropout. (D) Histograms of the resulting α from fits of the time-averaged ensemble MSDs using 500 bootstraps with 50% dropout. The resulting estimated parameters using 500 bootstraps with 50% dropout were $D^*_{x,y,z}$ = [1.10±0.02, 1.12±0.02, 1.04±0.02]$\times 10^{-3}$ $\mu m^2/s^{\alpha}$ and $\alpha_{x,y,z}$ = [0.437±0.005, 0.477±0.003, 0.407±0.007].

Supplemental Table S1. The measured anisotropy in $\alpha$ for untreated cells is independent of the z-correction factor used. Resulting mean D* and $\alpha$ from fits of the time-averaged ensemble MSD shown in Fig. 4 and Supplementary Figs. 4 and 10 for varying z-correction factors. The same experimental tracks were used for all calculations. D* increase with increasing correction factor, while $\alpha$ is unaffected by the correction. The corresponding resulting parameter values in x and y for untreated cells are: $D^*_{x,y}$ = [0.85, 0.83]$\times 10^{-3}$ $\mu m^2/s^\alpha$ and $\alpha_{x,y}$ = [0.38, 0.39], and for cells treated with LatB: $D^*_{x,y}$ = [1.10, 1.12]$\times 10^{-3}$ $\mu m^2/s^\alpha$ and $\alpha_{x,y}$ = [0.44, 0.48]. A correction factor of 0.75 results in roughly isotropic D* in all dimensions for cells treated with LatB. The anisotropy detected in $\alpha$ for untreated cells remains regardless of the chosen correction factor.

| | **Untreated cells** | | **Cells treated with LatB** | |
|---|---|---|---|---|
| **Correction factor** | **D*($\mu m^2/s^\alpha$)** | **$\alpha$** | **D*($\mu m^2/s^\alpha$)** | **$\alpha$** |
| **0.60** | $0.44\times 10^{-3}$ | 0.31 | $0.66\times 10^{-3}$ | 0.41 |
| **0.70** | $0.60\times 10^{-3}$ | 0.31 | $0.90\times 10^{-3}$ | 0.42 |
| **0.75** | $0.68\times 10^{-3}$ | 0.31 | $1.04\times 10^{-3}$ | 0.41 |
| **0.78** | $0.74\times 10^{-3}$ | 0.31 | $1.12\times 10^{-3}$ | 0.41 |
| **0.88** | $0.94\times 10^{-3}$ | 0.31 | $1.42\times 10^{-3}$ | 0.41 |
| **1.00** | $1.22\times 10^{-3}$ | 0.31 | $1.84\times 10^{-3}$ | 0.41 |
